# A genetic disorder reveals a hematopoietic stem cell regulatory network co-opted in leukemia

**DOI:** 10.1101/2021.12.09.471942

**Authors:** Richard A. Voit, Liming Tao, Fulong Yu, Liam D. Cato, Blake Cohen, Xiaotian Liao, Claudia Fiorini, Satish K. Nandakumar, Lara Wahlster, Kristian Teichert, Aviv Regev, Vijay G. Sankaran

## Abstract

The molecular regulation of human hematopoietic stem cell (HSC) maintenance is therapeutically important, but limitations in experimental systems and interspecies variation have constrained our knowledge of this process. Here, we have studied a rare genetic disorder due to *MECOM* haploinsufficiency, characterized by an early-onset absence of HSCs *in vivo*. By generating a faithful model of this disorder in primary human HSCs and coupling functional studies with integrative single-cell genomic analyses, we uncover a key transcriptional network involving hundreds of genes that is required for HSC maintenance. Through our analyses, we nominate cooperating transcriptional regulators and identify how MECOM prevents the CTCF-dependent genome reorganization that occurs as HSCs differentiate. Strikingly, we show that this transcriptional network is co-opted in high-risk leukemias, thereby enabling these cancers to acquire stem cell properties. Collectively, we illuminate a regulatory network necessary for HSC self-renewal through the study of a rare experiment of nature.

## INTRODUCTION

Human hematopoietic stem cells (HSCs) lie at the apex of the hierarchical process of hematopoiesis and rely on an intricate balance of transcriptional regulators to coordinate self-renewal and lineage commitment, and enable effective and continuous blood cell production^1^. Perturbations of HSC maintenance or differentiation result in a spectrum of hematopoietic consequences, ranging from bone marrow failure to leukemic transformation^2,3^. Despite the importance of HSCs in human health and the therapeutic opportunities that could arise from being able to better manipulate these cells, the precise regulatory networks that maintain these cells remain poorly understood.

Recently, loss-of-function mutations in Myelodysplastic Syndrome (MDS) and Ecotropic Virus Integration site-1 (EVI1) complex locus (*MECOM*) have been identified that lead to a severe neonatal bone marrow failure syndrome^4–15^. Strikingly, haploinsufficiency of this gene leads to near complete loss of HSCs within the first months of life, suggesting an important and dosage-dependent role for MECOM in early hematopoiesis. The role of MECOM in hematopoiesis has been studied using mouse models; homozygous *Evi1* knockout animals, which impact both the shorter *Evi1* and longer *Mecom* isoforms, are embryonic lethal and have pancytopenia with a paucity of HSCs^16–18^. Inducible knockout of *Evi1* in adult mice causes progressive pancytopenia and loss of HSCs, as well as downstream progenitors^17^. *Evi1* haploinsufficient mice display an intermediate phenotype, showing a reduced, but not absent, ability for hematopoietic reconstitution with maintenance of normal hematopoietic differentiation^17,19^. Endogenous disruption of the Mds-encoding region of one *Mecom* allele to generate a fluorescent reporter resulted in no observable defects in hematopoiesis^20^. These mouse studies reveal that different *Mecom* isoforms lead to varied functional consequences, but the ability of *Mecom* haploinsufficient mice to maintain sufficient hematopoietic output stands in sharp contrast to the profound and highly-penetrant HSC loss observed in patients with *MECOM* haploinsufficiency, irrespective of which isoform is impacted. These differences highlight interspecies variation in the role of MECOM in the maintenance of HSCs and suggest that these clinical observations may provide a unique experiment of nature to better understand human HSC regulation.

*MECOM* overexpression as a result of chromosome 3 aberrations has been reported in ∼10% of adult and pediatric acute myeloid leukemia (AML), and is associated with a particularly poor prognosis^21,22^. A number of mechanistic studies have highlighted specific targets of MECOM regulation in AML cell lines^23–27^. Despite the distinct potential mechanisms that have been suggested, the holistic functions of MECOM that enable effective human HSC maintenance remain enigmatic. Here, by taking advantage of *in vivo* observations from *MECOM* haploinsufficient patients, we have modeled this disorder through genome editing of primary human CD34^+^ hematopoietic stem and progenitor cells (HSPCs). Through integrative single-cell genomic analyses, we provide a refined understanding of the fundamental transcriptional regulatory circuits necessary for human HSC maintenance. Finally, we demonstrate that this same transcriptional regulatory network from human HSCs is co-opted in AML, thereby conferring stem cell features and a poor prognosis.

## RESULTS

### MECOM loss impairs HSC function *in vitro* and *in vivo*

Monoallelic mutations in *MECOM* have been implicated in severe, early onset neonatal aplastic anemia that is characterized by a paucity of hematopoietic cells. To date, at least 26 patients have been described with missense, nonsense, and frameshift mutations, as well as large deletions in *MECOM* that impact one or all isoforms (**Fig. 1a** **and Supplementary Table 1**). Nearly all of the missense mutations occur in exon 11 within a highly mutationally constrained zinc finger DNA-binding domain, where they are predicted to disrupt secondary structure and interfere with zinc coordination (**Extended Data Fig. 1a,b**).

**Figure 1.**
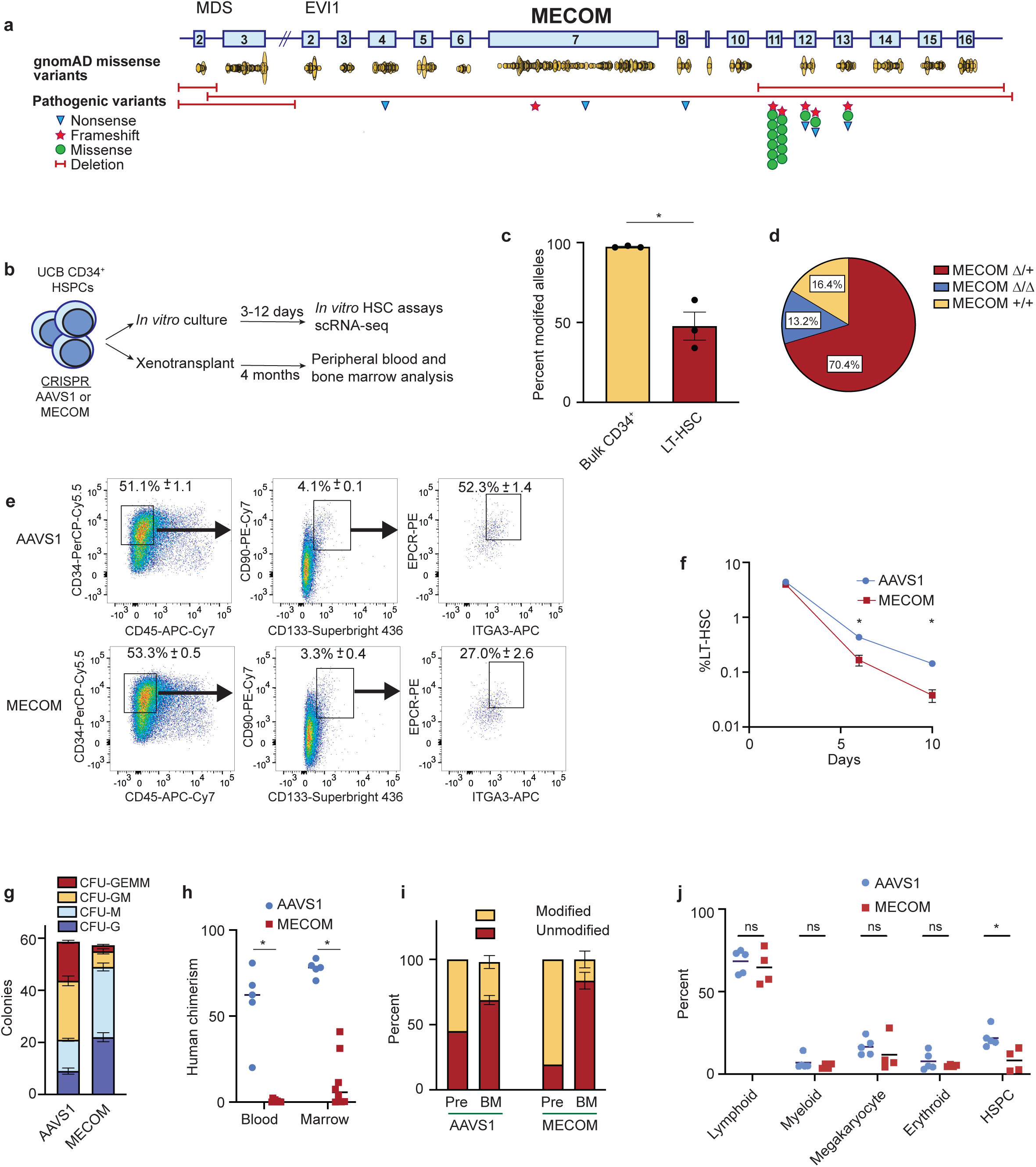
Generating a faithful model of *MECOM* haploinsufficiency and HSC loss. **(a)** Schematic of the *MECOM* locus displaying 2 coding exons of *MDS* (MDS 2-3) and 15 coding exons of *EVI1*(EVI1 2-16). Yellow ovals represent frequency and location of missense variants from individuals in the gnomAD database. Pathogenic variants from patients with bone marrow failure include nonsense (blue triangles), frameshift (red stars), and missense mutations (green circles) as well as large deletions (red bars). **(b)** Experimental outline of *MECOM* editing and downstream analysis in human umbilical cord blood-derived HSCs. **(c)** Bar graph of the frequency of modified *MECOM* alleles in bulk CD34^+^ human HSPCs or in LT-HSCs. HSPCs underwent CRISPR editing and were cultured in HSC media containing UM171. On day 6 after editing, genotyping by PCR and Sanger sequencing was performed on bulk HSPCs or LT-HSCs sorted by FACS. Mean of three independent experiments is plotted and error bars show s.e.m. Two-sided Student t-test used. \**P*<5e-3. **(d)** Pie chart showing the proportion of *MECOM* genotypes in single cell LT-HSCs following *MECOM* perturbation. 189 single cell LT-HSCs were genotyped using single cell genomic DNA sequencing and classified as either wild-type (MECOM^+/+^, yellow), heterozygous edited (MECOM^Δ/+^, red), or homozygous edited (MECOM^Δ/Δ^, blue). **(e-f)** Phenotypic analysis of LT-HSCs after *MECOM* editing. **(e)** Gating strategy to identify phenotypic LT-HSCs after CRISPR editing of *AAVS1* or *MECOM*. LT-HSCs are defined as CD34^+^CD45RA^-^CD90^+^CD133^+^EPCR^+^ITGA3^+^. Mean (± s.e.m.) in the highlighted gates on day 6 after CRISPR editing is shown (*n*=3), and the total LT-HSC percentage is the product of the frequencies in each gate shown. **(f)** Time course showing that *MECOM* editing leads to progressive loss of phenotypic LT-HSCs *in vitro.* X-axis displays days after CRISPR editing. Mean of three independent experiments is plotted and error bars show s.e.m. Two-sided Student *t*-test used. \**P* < 5e-3. **(g)** Stacked bar plots of colony forming assay comparing *MECOM* edited UCB-derived CD34^+^ HSPCs (*n*=3) to *AAVS1* edited controls (*n*=3). Three days after CRISPR perturbation, cells were plated in methylcellulose and colonies were counted after 14 days. *MECOM* editing leads to reduced formation of multipotent CFU-GEMM and bipotent CFU-GM progenitor colonies and an increase in unipotent colonies. CFU-GEMM, colony-forming unit (CFU) granulocyte erythroid macrophage megakaryocyte; CFU-GM, CFU granulocyte macrophage; CFU-M, CFU macrophage; CFU-G, CFU granulocyte. Mean colony number is plotted and error bars show s.e.m. **(h)** Analysis of peripheral blood and bone marrow of mice at week 16 following transplantation of *MECOM*-edited (*n*=8) and *AAVS1*-edited (*n*=4) HSPCs. Mean is indicated by black line and each data point represents one mouse. Two-sided Student *t*-test used. \**P* < 5e-6. **(i)** Comparison of edited allele frequency following xenotransplantation. *MECOM*-edited cells in bone marrow after xenotransplantation are enriched for unmodified alleles as detected by NGS, revealing a selective engraftment disadvantage of HSPCs with *MECOM* edits (Pre, pre-transplant; BM, bone marrow). Mean is plotted and error bars show s.e.m. **(j)** Subpopulation analysis of human cells in mouse bone marrow after xenotransplantation. Cell populations were identified by the following surface markers: lymphoid, CD45^+^CD19^+^; myeloid, CD45^+^CD11b^+^; megakaryocyte, CD45^+^CD41a^+^; erythroid, CD235a^+^; HSPC, CD34^+^. Only mice with human chimerism >2% were included in the analysis (AAVS1, 4/4 mice; MECOM, 4/8 mice). Mean is indicated by black lines and each data point represents one mouse. Two-sided Student *t*-test used. ns, not significant, \**P* = 0.01.

The profound bone marrow hypocellularity and absence of HSCs associated with *MECOM* haploinsufficiency prevents the mechanistic study of primary patient samples^5^. We therefore sought to develop a model to study *MECOM* haploinsufficiency in primary human cells by performing targeted disruption of *MECOM* via CRISPR editing in CD34^+^ HSPCs purified from umbilical cord blood (UCB) samples of healthy newborns (**Fig. 1b, Extended Data Fig. 1a,c,d**). We achieved editing at >80% of alleles in the bulk CD34^+^ population, but notably the subpopulation of phenotypic long-term HSCs (LT-HSCs)^28^ displayed 48% editing (**Fig. 1c**), consistent with the hypothesis that functional MECOM is crucial for the maintenance of LT-HSCs, and cells with MECOM perturbations more readily differentiate. Genotyping of individual single LT-HSCs following *MECOM* perturbation confirmed that 70% of LT-HSCs were heterozygous for *MECOM* edits (**Fig. 1d**), although this likely underestimates the true percentage of heterozygous edits given that allelic dropout is common in single cell genotyping^29^. These edits were faithfully transcribed to mRNA, but *MECOM* editing led to a significant reduction in *MECOM* mRNA levels in LT-HSCs, possibly due to nonsense-mediated decay^30^ (**Extended Data Fig. 1e-g**).

Prior work has shown that biallelic *Mecom* disruption in mouse HSCs results in a loss of quiescence, accompanied by increased cell cycle progression and differentiation^18^. Consistent with this observation, compared to *AAVS1*-edited cells, *MECOM*-edited human HSPCs underwent 1.9-fold higher expansion over 5 days in culture conditions that promote HSC maintenance (**Extended Data Fig. 1h,i**). *MECOM* perturbation was associated with a small but significant decrease in the proportion of bulk cells in G0/G1 on day 5 after CRISPR editing, but no difference in cell cycle states of HSCs (**Extended Data Fig. 1j**). Most HSCs remained in G0/G1 as analyzed by EdU incorporation and 7-AAD staining, and the majority of LT-HSCs had G0/G1 transcriptional signatures (**Extended Data Fig. 1k**), as previously reported^31^. *MECOM* editing resulted in more frequent cell divisions (**Extended Data Fig. 1l**) and a significant reduction in the absolute number of LT-HSCs **(Extended Data Fig. 1m)**. This resulted in a progressive loss of phenotypic LT-HSCs following *MECOM* editing with a 3.7-fold reduction by day 10 after editing (**Fig. 1e,f**). Together, these findings suggest that *MECOM* perturbation promotes the differentiation of LT-HSCs into more mature progenitors, which then expand while undergoing active progression through the cell cycle.

As further evidence that *MECOM* editing causes an impairment of HSCs *in vitro*, we observed a 6.4-fold reduction in multipotent CFU-GEMM colonies and a 3.8-fold reduction in bipotent CFU-GM colonies, along with increases in differentiated unipotential CFU-G and CFU-M colonies (**Fig. 1g**). There was a similar loss of multipotent and bipotent progenitor colonies derived from adult HSPCs following *MECOM* editing (**Extended Data Fig. 1n**), validating the importance of this factor across developmental stages.

Next, we performed non-irradiated transplantation of edited HSPCs into NBSGW mice to assess how MECOM loss impacts human HSCs *in vivo*^32–34^. Cells that underwent CRISPR editing of the control *AAVS1* locus engrafted well in all transplanted animals. In comparison, *MECOM*-edited cells engrafted in only half of the transplanted animals with significantly lower human chimerism in the peripheral blood and bone marrow (**Fig. 1h**). Since CRISPR editing results in a population of cells with heterogeneous genomic lesions, we compared the edited allele frequency of cells harvested from the bone marrow at 16 weeks with the cells prior to transplant and found a 5-fold enrichment of the unmodified *MECOM* allele (**Fig. 1i, Extended Data Fig. 1o,p**), consistent with selection occurring against HSCs that underwent editing at this locus. Having established that there is a significant reduction in total cell engraftment, we analyzed the bone marrow of the transplanted mice and found a 2.7-fold reduction in CD34^+^ HSPCs in the *MECOM*-edited samples, but no significant differences in engrafted lymphoid, erythroid, megakaryocytic, or monocytic lineages (**Fig. 1j**). Next, we evaluated engraftment of adult HSPCs following *MECOM* editing and found a comparable reduction in human chimerism in the bone marrow compared to *AAVS1* edited controls (**Extended Data Fig. 1q**). To evaluate serial repopulating ability of *MECOM*-perturbed UCB derived HSCs, we performed secondary xenotransplantation and observed moderate detectable secondary engraftment of *AAVS1* edited cells (2/5 mice), but no detectable secondary engraftment of *MECOM* edited cells (0/8 mice). To more sensitively assay for the presence of human cells in the secondary transplant recipients, we used human *MECOM*-specific PCR primers and amplified human *MECOM* from all bone marrow samples. Sequencing revealed 100% wild type *MECOM* in 7/8 secondary recipients and 95% in the remaining mouse (**Extended Data Fig. 1r**). This near complete absence of *MECOM* edits in serially-repopulating LT-HSCs is consistent with the profound HSC loss observed in patients with *MECOM* haploinsufficiency.

In sum, our model of *MECOM* haploinsufficiency reveals that MECOM is required for maintenance of LT-HSC *in vitro* and *in vivo*. Crucially, this model enables us to create *MECOM* haploinsufficiency and capture LT-HSCs prior to their complete loss, thus allowing for the direct study of the MECOM function in primary human LT-HSCs.

### MECOM loss in LT-HSCs elucidates a dysregulated gene network

Having confirmed that our model of *MECOM* loss in primary human CD34^+^ HSPCs faithfully recapitulates the profound HSC defect observed in patients with *MECOM* haploinsufficiency, we sought to use this system to gain mechanistic insights into the transcriptional circuitry required for human HSC maintenance by single-cell RNA sequencing (scRNA-seq) prior to complete HSC loss. We performed CRISPR editing of *MECOM* in UCB CD34^+^ HSPCs and maintained the cells in HSC media containing UM171 for 3 additional days prior to sorting for phenotypic LT-HSCs and performing scRNA-seq using the 10x Genomics platform. We reasoned that scRNA-seq in this sorted compartment was necessary, given the known heterogeneity present among HSCs^1,35^, as well as the heterogeneous editing outcomes that would occur. To confirm the fidelity of our sorting strategy, we examined the expression of an HSC signature (*CD34*, *HLF*, *CRHBP*)^36^, which is found in a rare subpopulation representing only 0.6% of 263,828 cord blood cells from the immune cell atlas (**Fig. 2a**), and observed that our sorted phenotypic LT-HSCs are highly enriched for this HSC signature (**Fig. 2b, Extended Data Fig. 2a-c**). Next, we compared the transcriptomes of 5,935 *MECOM*-edited and 4,291 *AAVS1*-edited phenotypic LT-HSCs. *MECOM*-edited LT-HSCs co-localized with *AAVS1*-edited cells in uniform manifold approximation and projection (UMAP) space (**Fig. 2c, Extended Data Fig. 2d,e**), confirming that our sorting strategy would allow us to directly compare developmentally stage-matched cells, and that these cells share high-dimensional transcriptional similarity.

**Figure 2.**
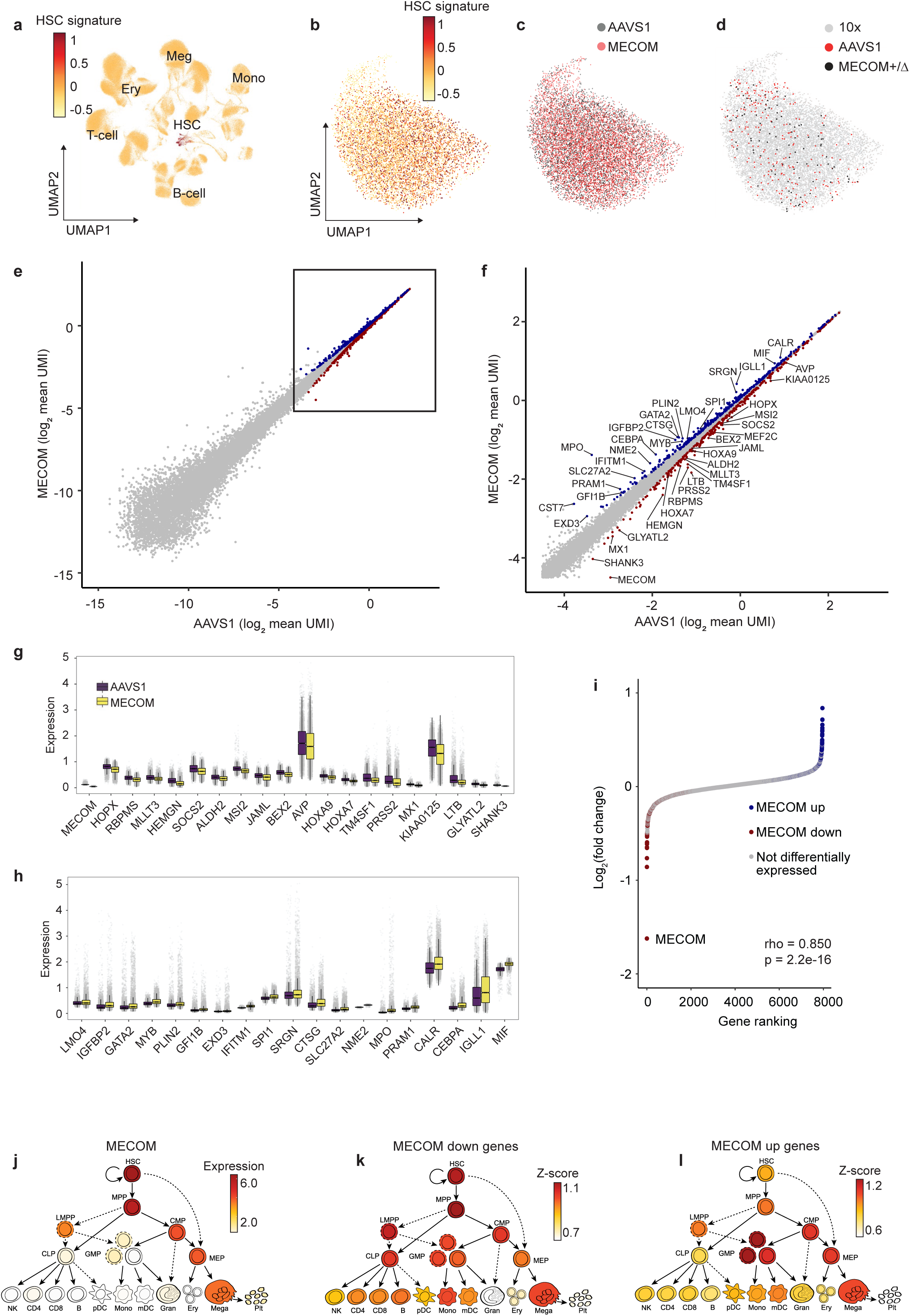
Delineation of a MECOM regulatory network in LT-HSCs. **(a)** Uniform manifold approximation and projection (UMAP) plot of 263,828 single cells from human umbilical cord blood, colored according to HSC signature (*CD34*, *HLF*, *CRHBP*). **(b-d)** UMAP plots of phenotypic LT-HSCs following CRISPR editing, indicating **(b)** enrichment of the HSC signature as determined by scRNA-seq using the 10x Genomics platform, **(c)** overlap of *AAVS1* edited and *MECOM* edited cells, sequenced using the 10x Genomics platform, and **(d)** distribution of cells with monoallelic *MECOM* edits determined by G&T sequencing by Smart-Seq2, compared to *AAVS1* edited cells, and LT-HSCs from **(b)** and **(c)**. **(e-f)** Scatter plots of gene expression in LT-HSCs following *AAVS1* or *MECOM* editing. Single cell expression data for each gene was averaged following imputation, and is plotted. Differential gene expression was determined using Seurat 4.0 differential expression analysis with the MAST pipeline, and is indicated by colored dots, MECOM down genes, red; MECOM up genes, blue. **(e)** displays the expression of all genes, and **(f)** displays a subset containing the most highly expressed genes. A gene is defined as differentially expressed if the log_2_ fold change is greater than 0.05 and the adjusted p-value is less than 1e-20. **(g-h)** Box plots showing expression of a subset of MECOM down **(g)** and MECOM up **(h)** genes after *MECOM* editing. Gray dots show imputed gene expression in single cells. **(i)** Pseudobulk analysis of differentially expressed genes. Transcriptomic data from single LT-HSCs that had undergone *AAVS1* or *MECOM* perturbation were integrated to generate pseudobulk gene expression profiles. Expression differences between the AAVS1 and MECOM pseudobulk samples are plotted in rank order, and differentially expressed genes from the scRNA-seq analysis are highlighted (MECOM down genes, red; MECOM up genes, blue). Correlation of differential gene expression between pseudobulk and single cell analyses was calculated using Spearman’s rank correlation. **(j-l)** Expression of *MECOM* (log_2_ normalized CPM) throughout hematopoietic differentiation reveals robust expression in HSCs **(j)**, similar to the enrichment of expression of MECOM down genes **(k)** and the inverse of the expression pattern of MECOM up genes **(l)**.

As an orthogonal approach to simultaneously profile the precise genomic editing outcome and transcriptional profile of LT-HSCs, we employed genome and transcriptome sequencing (G&T-seq). *MECOM* heterozygous cells (**Fig. 1d**) colocalize in UMAP space with *AAVS1* edited cells, as well as both the non-genotyped cells examined with the 10X Genomics method (**Fig. 2d**). These results reveal a high degree of similarity in the high-dimensional transcriptomic analysis of LT-HSCs following *MECOM* perturbation, as expected given the stringent phenotypic sorting strategy we employed prior to single cell RNAs sequencing analysis. Furthermore, these results suggest that the profound functional consequences of MECOM loss are due to coordinated expression changes in a select group of genes.

To compare individual gene expression in single LT-HSCs following *AAVS1* or *MECOM* editing, we used model-based analysis of single-cell transcriptomes (MAST)^37^ (**Fig 2e,f, Extended Data Fig. 2f**). Despite the high-dimensional transcriptional similarity, we detected significant downregulation of a group of 322 genes following *MECOM* editing that we refer to as ‘MECOM down’ genes (**Supplementary Table 2**). Not surprisingly, the MECOM down gene set includes factors necessary for HSC self-renewal and maintenance including *HOPX*, *HOXA9*, *RBPMS*, *MLLT3*, *MEF2C*, *HEMGN*, *SOCS2*, *ALDH2*, *HLF*, *MSI2*, *ALDH1A1*, and *ADH5* (**Fig. 2f,g**), but also uncovers genes without known regulatory functions in HSCs. We then used MAST to identify 402 genes that are significantly upregulated after *MECOM* editing, which we refer to as the ‘MECOM up’ gene set (**Supplementary Table 2**). The MECOM up gene set includes key factors expressed during hematopoietic differentiation including *MPO*, *SPI1*, *CALR*, *CEBPA*, *MIF*, *GATA2*, and *GFI1B* (**Fig. 2f,h**), but also identifies genes without known roles in hematopoietic differentiation or lineage commitment. To validate that the MECOM down and up gene sets represented true biological differences rather than random stochastic variation, we performed permutation analysis and did not detect a single significant differentially expressed gene in random permutations, including no differential expression of any MECOM down or MECOM up genes, highlighting the robustness of our differential gene analysis (**Extended Data Fig. 2g,h**).

Additionally, to minimize the potential confounding influence of allelic dropout, we performed pseudobulk analysis of gene expression changes following *MECOM* perturbation^38^. We observed that the nominated MECOM down and up gene sets again represented the most differentially expressed genes with larger expression differences compared to the single cell analysis (**Fig. 2i**). To validate that the gene expression differences that we observed in the population of phenotypic LT-HSCs accurately represented gene expression changes in transcriptional LT-HSCs, we examined expression of each differentially expressed gene in the subset of phenotypic LT-HSCs with robust expression of the HSC signature (**Fig. 2b**). There was significant correlation of gene expression changes in this subpopulation of transcriptional LT-HSCs compared to the bulk cells, demonstrating that MECOM network genes were indeed differentially expressed in cells with a stringent molecular HSC signature (**Extended Data Fig. 2i**).

We then evaluated the expression of the MECOM down and up genes during normal hematopoiesis by comparing the enrichment of the gene sets in 20 distinct hematopoietic cell lineages^39^. Similar to the expression pattern of *MECOM* itself (**Fig. 2j**), the MECOM down genes are collectively more highly expressed in HSCs and earlier progenitors compared to more differentiated cells (**Fig. 2k**). Conversely, the MECOM up genes are turned on during hematopoietic differentiation (**Fig. 2l**). Collectively, these analyses reveal that MECOM loss in LT-HSCs leads to functionally significant transcriptional dysregulation in genes that are fundamental to HSC maintenance and differentiation.

### Increased *MECOM* expression rescues functional and transcriptional changes in HSCs

To confirm that the functional and transcriptional impacts on LT-HSCs that we observed are due specifically to reduced MECOM levels, we sought to rescue the phenotype by lentiviral *MECOM* expression in HSCs after CRISPR editing (**Fig. 3a**). To avoid unintended CRISPR disruption of the lentivirally encoded *MECOM* cDNA, we introduced wobble mutations in the sgRNA binding site in the cDNA (**Extended Data Fig. 3a,b**). Infection of *MECOM*-edited HSPCs with *MECOM* encoding virus led to supraphysiologic levels of *MECOM* expression (**Fig. 3b**). Functionally, this *MECOM* overexpression was sufficient to rescue the LT-HSC loss observed after *MECOM* editing and resulted in preservation of more LT-HSCs compared to control samples on day 6 after CRISPR editing (**Fig. 3c,d, Extended Data Fig. 3c,d**). We also examined the ability of other MECOM isoforms to expand LT-HSCs in *AAVS1*- and *MECOM*-edited HSPCs. Increased expression of *EVI1* resulted in a higher percentage of LT-HSCs on day 6 in culture, but this increase was blunted by endogenous *MECOM* editing. Expression of *MDS* did not result in rescue of LT-HSCs (**Extended Data Fig. 3e**). Together, these data reveal that restoration of the full length *MECOM* isoform is sufficient to overcome the functional loss of LT-HSCs caused by endogenous *MECOM* perturbation. Since the *MECOM* virus co-expresses GFP, we reasoned that cells that remained in the LT-HSC subpopulation after *MECOM* editing and infection would be enriched for increased *MECOM* expression and therefore GFP expression. Indeed, in samples transduced with the *MECOM* virus, we observed a significantly higher ratio of GFP expression in LT-HSCs compared to the bulk population (**Fig. 3e**). Increased *MECOM* expression was also sufficient to rescue the loss of multipotent and bipotent progenitor colonies after *MECOM* editing (**Fig. 3f**).

**Figure 3.**
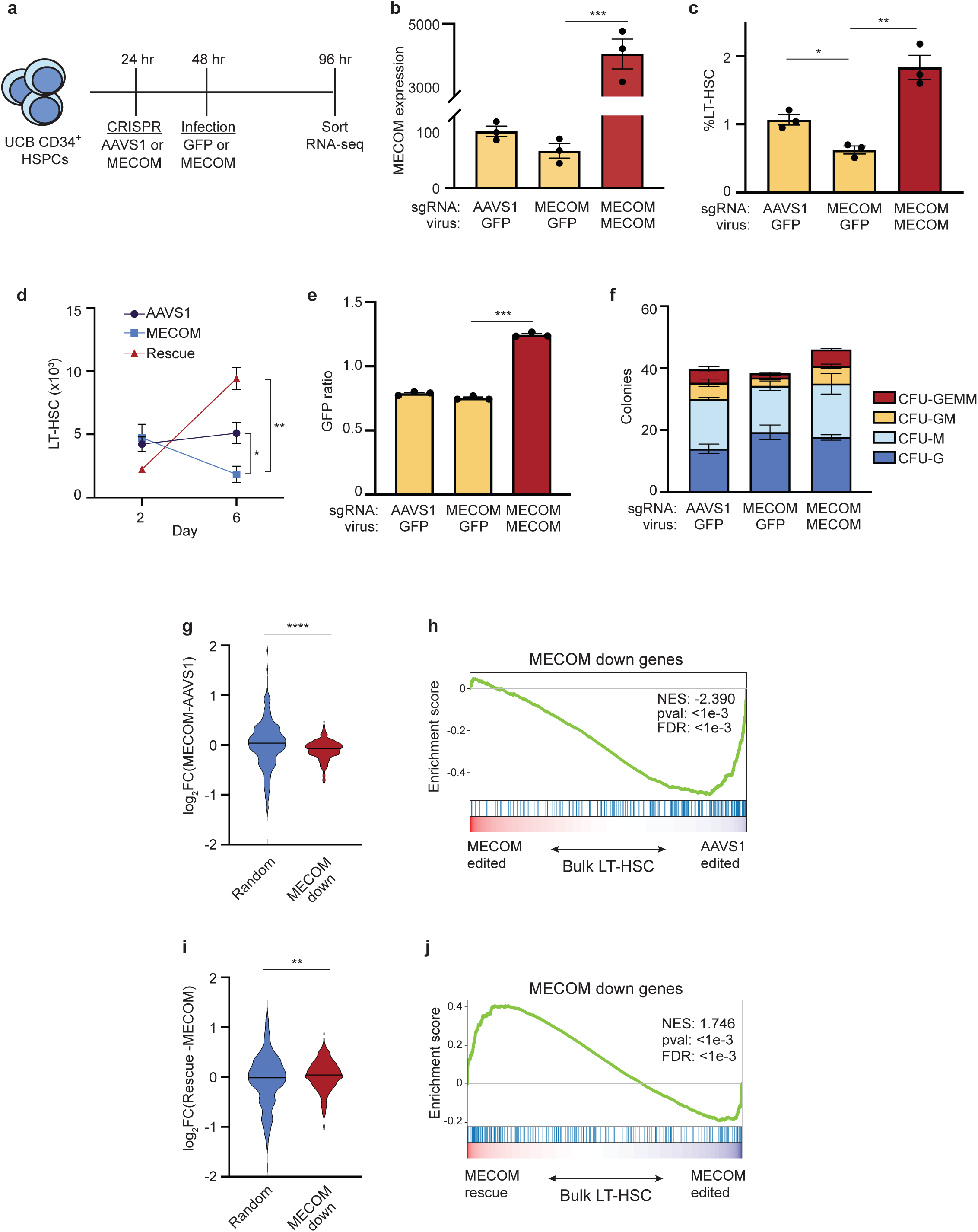
MECOM rescue of functional and transcriptional changes in HSCs. **(a)** Experimental outline of *MECOM* editing and rescue. **(b-d)** Effects of *MECOM* editing and infection with *MECOM* or GFP lentivirus. **(b)** *MECOM* expression (RPKM) measured by RNA-seq is shown. **(c)** Percent of LT-HSC determined by FACS, and **(d)** number of LT-HSCs are shown. *n*=3 per group. Mean is plotted and error bars show s.e.m. Two-sided Student *t*-test used. \**P* < 5e-2, \*\**P* < 5e-3, \*\*\**P* < 5e-4. **(e)** GFP ratio following lentiviral infection. GFP ratio is defined as percent of GFP^+^ LT-HSCs divided by the percent GFP^+^ bulk HSPCs. GFP ratio >1 is consistent with enrichment of infected cells in the LT-HSC population. *n*=3 per group. Mean is plotted and error bars show s.e.m. Two-sided Student *t*-test used. \*\*\**P* < 5e-4. **(f)** Stacked bar plots of colony forming assay. Infection with *MECOM* virus leads to restoration of multipotent CFU-GEMM and bipotent CFU-GM colonies that are lost following MECOM editing, *n*=3 per group. CFU-GEMM, colony-forming unit (CFU) granulocyte erythroid macrophage megakaryocyte; CFU-GM, CFU granulocyte macrophage; CFU-M, CFU macrophage; CFU-G, CFU granulocyte. Mean colony number is plotted and error bars show s.e.m. **(g)** Violin plot of differential gene expression in bulk LT-HSCs following *MECOM* perturbation. MECOM down genes are significantly depleted in *MECOM* edited samples compared to AAVS1 edited samples, unlike a set of randomly selected genes. Two-sided Student *t*-test used. **** *P* < 1e-4. **(h)** GSEA of MECOM down genes after *MECOM* perturbation. MECOM down genes that were identified from the single cell RNA sequencing analysis are depleted in *MECOM* edited LT-HSCs in bulk, compared to *AAVS1* edited cells. **(i)** Violin plot of differential gene expression in bulk LT-HSCs following *MECOM* perturbation and rescue. MECOM down genes are significantly enriched in *MECOM* rescue samples compared to *MECOM* edited samples, unlike a set of randomly selected genes. Two-sided Student *t*-test used. ** *P* < 5e-3. **(j)** GSEA of MECOM down genes after *MECOM* perturbation and rescue. MECOM down genes that were identified from the single cell RNA sequencing analysis are enriched in *MECOM* rescued LT-HSCs in bulk, compared to *MECOM* edited cells.

Next, we examined the transcriptional profile of phenotypic LT-HSCs after *MECOM* editing and rescue. We performed CRISPR editing of UCB-derived CD34^+^ HSPCs, followed by *MECOM* or GFP virus infection. Cells were sorted for expression of GFP and phenotypic LT-HSC markers on day 4 and subjected to RNA sequencing. Following *MECOM* perturbation alone, we observed significantly lower expression of the MECOM down gene set compared to a subset of randomly selected genes, as expected (**Fig. 3g**). Similarly, GSEA analysis revealed significant depletion of the MECOM down genes (**Fig. 3h**). Following increased *MECOM* expression, the MECOM down genes were significantly upregulated (**Fig. 3i,j, Supplementary Table 3**). Interestingly, we did not observe similar upregulation or subsequent rescue of the MECOM up genes in bulk following *MECOM* perturbation and overexpression (**Extended Data Fig. 3f,g**). This may be attributable to the different temporal pattern of expression between the gene sets; MECOM down genes are expressed in HSCs and are necessary for maintenance, while the MECOM up genes come on during differentiation. Alternatively, the supraphysiologic expression that we obtained may not allow effective regulation of the MECOM up genes. Regardless, these data collectively show that the loss of LT-HSCs after *MECOM* editing can be restored with increased *MECOM* expression and is accompanied by the rescue of the MECOM down gene set.

### Defining the HSC *cis*-regulatory network mediated by MECOM

Having identified a set of dysregulated genes after loss of *MECOM* in LT-HSCs, we next sought to define the *cis*-regulatory elements (*cis*REs) that control expression of this MECOM dependent gene network that underlies HSC self-renewal. To do so, we developed HemeMap, a computational framework to identify putative *cis*REs and cell type-specific *cis*RE-gene interactions by integrating multi-omic data from 18 cell populations across distinct hematopoietic lineages (**Fig. 4a, Extended Data Fig. 4a,b**)^40–44^, and calculated a HemeMap score based on the chromatin accessibility for each *cis*RE-gene interaction in HSCs. We found that the HemeMap scores were closely correlated with gene expression (**Extended Data Fig. 4c**). All of the interactions with a significant HemeMap score in HSCs were selected to construct an HSC-specific regulatory network (**Extended Data Fig. 4d**).

**Figure 4.**
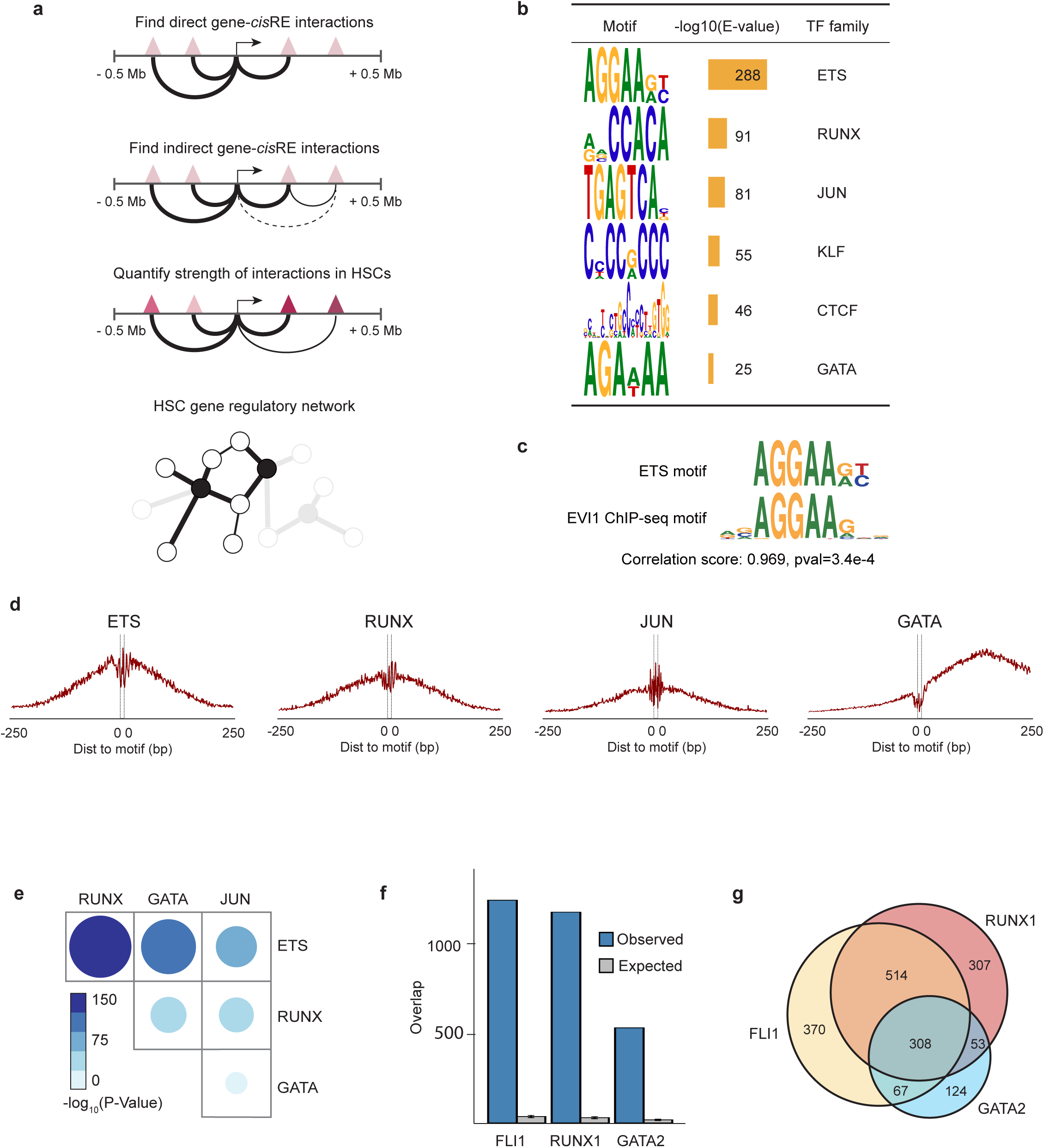
Defining the HSC *cis*-regulatory network coordinated by MECOM. **(a)** Schematic of the HemeMap method used to define an HSC-specific regulatory network. **(b)** Significantly enriched conserved motifs associated with *cis*REs of MECOM network genes in the HSC-specific regulatory network. Motif discovery and significance testing were performed using MEME. **(c)** Motif similarity between the ETS motif and a previously identified EVI1 motif from ChIP-seq^24^. Similarity was determined by the Pearson correlation coefficient of the Position Frequency Matrix in a comparison of the two motifs. **(d)** Footprinting analysis of ETS, RUNX, JUN, and GATA within the *cis*REs in the MECOM regulation network. The plots show Tn5 enzyme cleavage probability of each base flanking (± 250 bp) and within TF motifs in HSCs. **(e)** Analysis of TF footprint co-occurrence in the MECOM network. The frequency of occurrence of each footprint in MECOM network *cis*REs was computed and the *P* value of co-occurrence for each TF pair was determined by a hypergeometric test. The color and size of dots are proportional to statistical significance. **(f)** Specific TF occupancy of *cis*REs in the MECOM network in CD34^+^ HSPCs. The number of *cis*REs associated with the MECOM network that overlap with ChIP-seq peaks for FLI1, RUNX1, and GATA2 were determined. For each TF, the expected distribution of overlapping *cis*REs was generated by 1,000 permutations of an equal number of TF peaks across the genome. Mean is plotted and error bars show s.d. **(g)** Overlap of TF occupancy in MECOM network *cis*REs. The number of *cis*REs that contain ChIP-seq peaks for FLI1 (yellow), RUNX1 (red), GATA2 (blue) or combinations of TFs are indicated.

To identify the transcription factors (TFs) driving expression of the MECOM network genes, we performed unbiased motif discovery within the *cis*REs that we found to be associated with MECOM network genes in HSCs. We found six significantly enriched motifs: ETS, RUNX, JUN, KLF, CTCF, and GATA (**Fig. 4b**). The ETS family motif (AGGAAGT) was the most enriched TF binding motif in the *cis*REs of MECOM network genes and is a known binding site for several TFs that are thought to play a role in HSCs, including FLI1, ERG, ETV2, and ETV6^45^. Additionally and importantly, the experimentally-determined binding motif of EVI1 in AML cells^24^, is a near perfect mimic of our nominated ETS motif, suggesting that many of these sites may be directly occupied by MECOM (**Fig. 4c**). Highlighting the importance of ETS family members in the regulation of MECOM network genes, the HemeMap scores were significantly higher in *cis*REs with ETS motifs compared to those without (**Extended Data Fig. 4e**).

Next, we performed digital genomic footprinting analysis to filter the consensus motif sites and predict TF occupancy in HSCs (**Supplementary Tables 4,5**) and observed a clear pattern of TF occupancy in HSCs in the vicinity of the nominated footprints (**Fig. 4d**). We observed a significant co-occurrence of footprints across different TF pairs, with a particular enrichment of overlap of ETS motifs with RUNX, JUN, and GATA, suggesting cooperativity of these TFs. This also further emphasizes the central role of the ETS motif, which may be occupied by MECOM or other cooperating TFs, during the regulation of this functionally important HSC *cis*RE network (**Fig. 4e, Extended Data Fig. 4f,g**).

Next, we evaluated specific TF binding to the HSC *cis*REs nominated by the HemeMap analysis by integrating TF ChIP-seq data from human HSPCs^46^. Consistent with footprinting analysis of putative HSC TFs at the *cis*REs, we found highly enriched TF occupancy of the ETS family member FLI1, as well as RUNX1 and GATA2 (**Fig. 4f**) in HSPCs. Notably, these ChIP-seq data are derived from binding in bulk CD34^+^ HSPCs, so while they provide a general indication of TF binding in HSPCs, there may be important differences in TF binding in the rare subset of quiescent LT-HSCs. As further evidence of TF cooperativity, we found that FLI1, RUNX1, and GATA2 have striking co-occupancy at the MECOM-regulated gene *cis*REs in HSPCs (**Fig. 4g**). Together, these results suggest cooperativity among a number of key regulatory transcription factors that assist MECOM in regulating expression of MECOM network genes to enable effective HSC maintenance, and that may take the place of MECOM as cells differentiate from the HSC compartment.

### Dynamic CTCF binding during HSC activation represses MECOM down genes

In addition to the enrichment of important HSC transcription factor motifs, the *cis*REs of the MECOM gene network showed striking CTCF binding motif enrichment. CTCF is a key regulator of 3-dimensional genome organization and acts by both anchoring cohesin-based chromatin loops to insulate genomic regions of self-interaction, known as topologically associating domains (TADs), and by enabling looping between interacting regulatory elements^47–49^. Spatial orientation of neighboring motifs is crucial to the function of CTCF, and TAD boundaries are marked by divergent CTCF motifs. Within TADs, convergent CTCF sites co-occur with motifs of lineage-defining TFs and mediate *cis*RE-promoter interactions^50^. Recently, CTCF has been implicated in regulating HSC differentiation by altering looping and helping to silence key stemness genes^51^, while also cooperating with lineage-specific TFs during hematopoietic differentiation^52^. Therefore, we hypothesized that CTCF plays a role in mediating the differential expression of MECOM down genes following loss of *MECOM*. We focused these analyses on the MECOM down gene set since their expression is directly dependent on *MECOM* expression and necessary for HSC self-renewal, as shown through the *MECOM* rescue studies (**Fig. 3i,j**).

Footprinting analysis revealed high confidence CTCF footprints in bulk CD34^+^ HSPCs (**Fig. 5a**). There was moderate but significant co-occurence of CTCF footprint with ETS, RUNX, JUN, and KLF footprints in the *cis*REs of MECOM down genes (**Fig. 5b**). We observed a high level of CTCF binding to the nominated *cis*REs (**Fig. 5c**). Next, we compared CTCF binding in CD34^+^ HSPCs with terminally differentiated hematopoietic cells. We found CTCF occupancy of the nominated CTCF footprints was highly conserved across erythroid cells, T-cells, B-cells, and monocytes (**Fig. 5d, Extended Data Fig. 5a**). Notably, CTCF binding in HSPCs was measured in the population of bulk CD34^+^ cells, which contains, but is not limited to LT-HSCs. Despite this heterogeneity of the HSPC compartment, terminally differentiated cells showed significantly stronger CTCF signals compared to the CD34^+^ HSPCs and chromatin accessibility at those loci decreased during hematopoietic differentiation (**Extended Data Fig. 5b-d**). These results reveal increased binding of CTCF to the *cis*REs of MECOM down genes following HSC differentiation.

**Figure 5.**
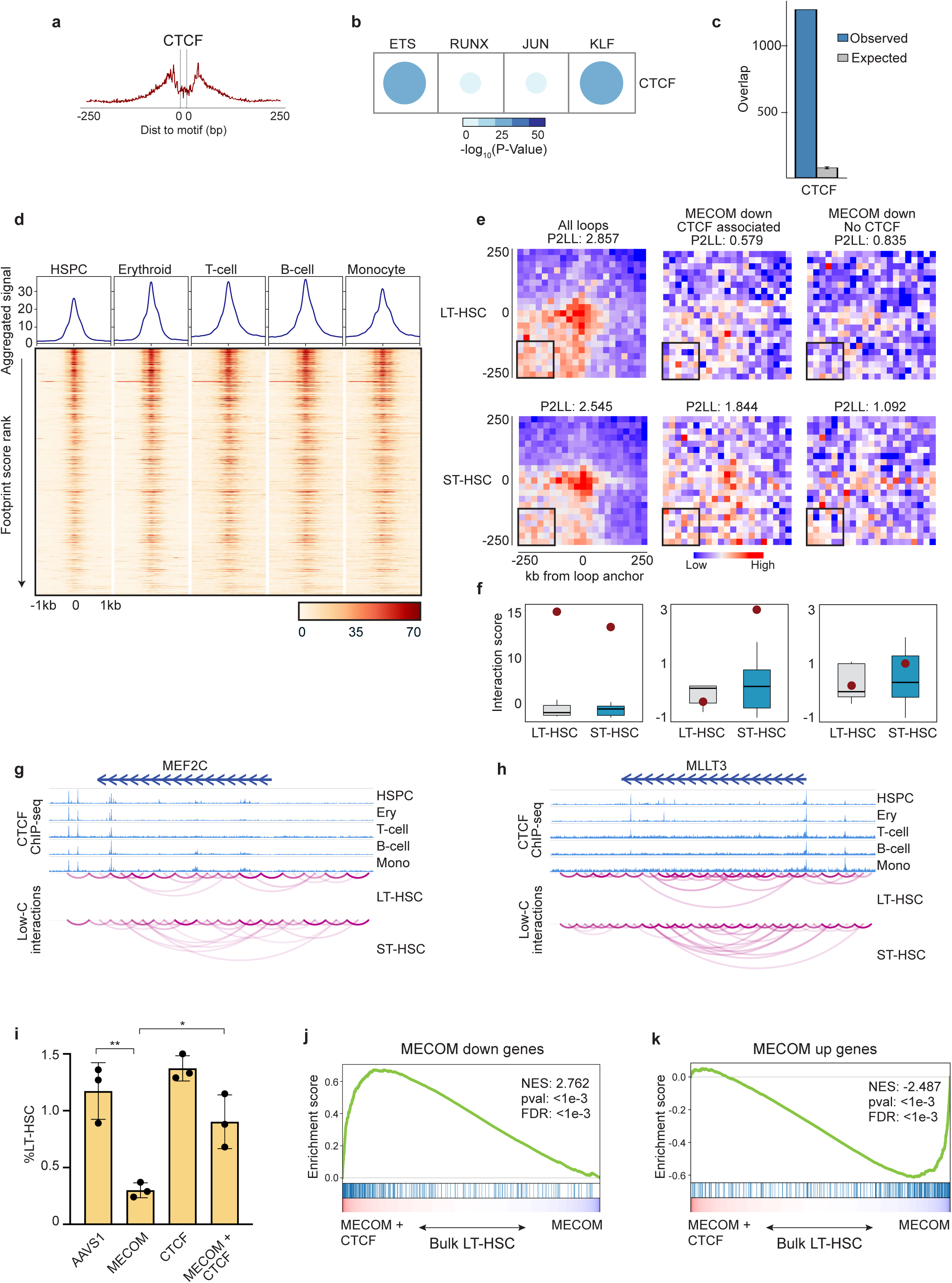
Dynamic CTCF binding facilitates repression of MECOM down genes as HSCs undergo differentiation. **(a)** Footprinting analysis of CTCF within the *cis*REs in the MECOM gene network. The plot shows Tn5 enzyme cleavage probability for each base flanking (± 250 bp) and within the CTCF motif. **(b)** Analysis of TF footprint co-occurrence of CTCF and other TFs in *cis*REs associated with MECOM down genes. The frequency of occurrence and *P* values were calculated using a hypergeometric test. The color and size of dots are proportional to statistical significance. **(c)** CTCF occupancy of *cis*REs in MECOM down genes in CD34^+^ HSPCs. The number of *cis*REs associated with the MECOM down genes that overlap with CTCF ChIP-seq peaks was determined and plotted as in Fig. 4f. **(d)** CTCF binding to MECOM down *cis*REs in hematopoietic lineages. Heatmaps (bottom) show the CTCF ChIP-seq signals that overlap CTCF footprints in MECOM down *cis*REs in HSPCs, erythroid cells, T-cells, B-cells, and monocytes. Each row represents a footprint ±1 kb of flanking regions, and the rows are sorted by the posterior probability of footprint occupancy from high to low. The enrichment of CTCF binding to *cis*REs was calculated and displayed in the line graph (top). **(e)** Aggregate peak analysis for the enrichment of chromatin loops in LT-HSCs (top) and ST-HSCs (bottom) using Low-C data. Chromatin loop interactions were determined for all chromatin loops derived from Hi-C data in hematopoiesis (left), the subset of CTCF-associated loops of MECOM down genes (center), and the subset of non-CTCF-associated loops of MECOM down genes (right). Aggregate signals over 500 kb centered on loop anchors with 25 kb resolution were calculated and are shown. The Peak to lower-left ratio (P2LL) enrichment score was calculated by comparing the peak signal to the mean signal of bins highlighted in black box in the heatmap and is shown in the title of each plot. **(f)** The standard normalized distribution of interaction scores for the lower left corner highlighted in the heatmap (Fig. 5e) is shown in the boxplots. Red dots indicate the peak value. The columns are as described in Fig. 5e. **(g-h)** Genome browser views of CTCF occupancy and chromatin interaction at *MEF2C* **(g)** and *MLLT3* **(h)** gene loci in LT-HSCs and ST-HSCs. **(i)** Bar graphs of LT-HSC rescue by dual *MECOM* and *CTCF* perturbation. Human HSPCs underwent CRISPR editing with the sgRNA guides depicted on the x-axis. Percent of LT-HSCs was determined by FACS on day 6. *n*=3 per group. Mean is plotted and error bars show s.e.m. Two-sided Student *t*-test used. \**P* < 1e-2, \*\**P* < 5e-3. **(j-k)** GSEA of MECOM down genes **(j)** and MECOM up genes **(k)** after dual *MECOM* and *CTCF* perturbation compared to *MECOM* perturbation alone. Bulk RNA sequencing was performed in biological triplicate on day 5 after CRISPR perturbation. MECOM down genes are enriched and MECOM up genes are depleted following concurrent *CTCF* editing.

To gain mechanistic insights into the role of CTCF in the MECOM-driven regulation of HSC quiescence, we analyzed an overall set of 7,358 chromatin loops from studies of HSCs^51^, as well as a subset of loops whose anchors co-localized with *cis*REs in the MECOM network. In total, 448 chromatin interactions were identified for MECOM down genes, and the loop anchors showed a strong enrichment of CTCF footprints (**Extended Data Fig. 5e**). Next, we performed aggregate peak analysis (APA) to compare the genomic organization of the MECOM down genes upon early exit from quiescence by integrating Low-C chromatin interaction data from phenotypic LT-HSCs and ST-HSCs^51^. Using all 7,358 common chromatin loops, there was significant enrichment of chromatin interaction apices in both LT-HSCs and ST-HSCs, as previously observed^51^, but there was no significant difference between LT-HSCs and ST-HSCs. Notably, analysis of the chromatin loops of CTCF footprint-containing *cis*REs associated with MECOM down genes revealed significantly stronger chromatin interactions in ST-HSCs compared to LT-HSCs. Importantly, there was no chromatin interaction difference in MECOM down genes that lacked association with a CTCF footprint-containing *cis*RE (**Fig. 5e,f**). These observations are consistent with the concept that CTCF binding to the *cis*REs of MECOM down genes induces tighter chromatin looping and restricted gene expression, promoting differentiation of HSCs, as exemplified by the increased chromatin looping at *MLLT3* and *MEF2C* concordant with their silencing during differentiation of LT-HSCs (**Fig. 5g,h**).

shRNA-mediated knockdown of CTCF in LT-HSCs prevents their exit from quiescence and induces transcriptional changes consistent with the maintenance of stemness^51^. Because of the correlation of the repression of MECOM down genes upon HSC activation by chromatin looping mediated by CTCF, we hypothesized that *CTCF* perturbation would lead to increased expression of MECOM down genes. We performed simultaneous *MECOM* and *CTCF* CRISPR perturbation in primary human UCB HSPCs (**Extended Data Fig. 5f**), and observed that concurrent *CTCF* perturbation was sufficient to rescue the loss of LT-HSCs induced by *MECOM* editing (**Fig. 5i**). Additionally, CTCF loss prevented the increased expansion of HSPCs caused by *MECOM* perturbation (**Extended Data Fig. 5g**).

Next, we examined the transcriptional changes that occur following dual *MECOM* and *CTCF* editing in LT-HSCs by RNA sequencing. First, we compared gene expression changes following *AAVS1* editing or *MECOM* editing alone. Using GSEA, we observed significant depletion of MECOM down genes and significant upregulation of MECOM up genes following *MECOM* editing alone, corroborating our observations from single cells (**Extended Data Fig. 5h,i**). Importantly, dual editing of *MECOM* and *CTCF* resulted in significant upregulation of MECOM down genes (**Fig. 5j**) and significant depletion of MECOM up genes (**Fig. 5k**). Upon dual perturbation, there was significantly greater rescue of MECOM down genes that are associated with *cis*REs containing CTCF binding motifs compared to those without CTCF motifs (**Extended Data Fig. 5j,k**). These data demonstrate that MECOM plays a key role in activating the expression of genes critical for HSC maintenance, which are then subject to genomic reorganization by CTCF as these cells undergo differentiation.

### The MECOM gene network is hijacked in high-risk AMLs

Having elucidated a fundamental transcriptional regulatory network necessary for HSC maintenance, we wondered to what extent this network may be relevant to leukemogenic states given the well-known role for *MECOM* overexpression in high-risk forms of AML. We reasoned that the transcriptional changes in MECOM network genes in LT-HSCs that we detected via sensitive single-cell RNA sequencing approaches might identify a transcriptional signature with prognostic implications in AML.

First, we combined 165 primary adult AML samples from The Cancer Genome Atlas (TCGA)^53^ with 430 adult samples from the BEAT AML dataset^54^ into an adult AML cohort (**Fig. 6a**) which we analyzed in parallel with 440 pediatric AML samples from the TARGET AML dataset^55^ (**Fig. 6b**). Prior reports from large cohorts of AML patients revealed a significant survival disadvantage in *MECOM*-high AMLs^56,57^. Using optimal thresholding to stratify patients by *MECOM* expression, we observed a similar poor prognosis in both the adult and pediatric datasets (**Fig. 6c**).

**Figure 6.**
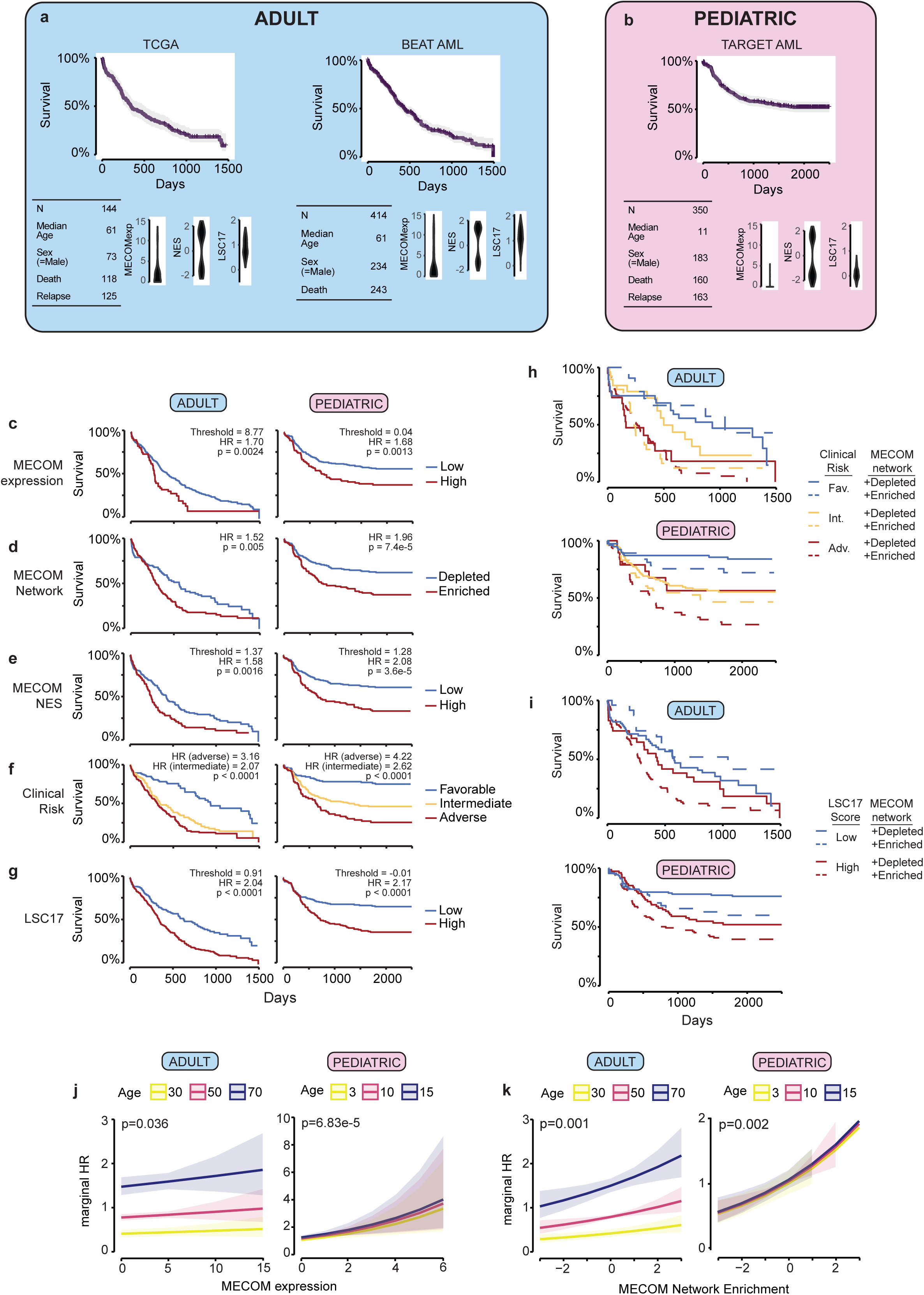
The MECOM down gene network is hijacked in high-risk adult and pediatric AML. **(a-b)** Descriptive statistics for included clinical cohorts. After correcting for study, TCGA and BEAT data were integrated into an adult cohort **(a)**. All of the pediatric data came from the TARGET database **(b)**. Distribution of MECOM expression, MECOM Network Enrichment Score (NES), and LSC17 score are displayed for each clinical dataset. **(c-g)** Kaplan-Meier (KM) overall survival curves for adult and pediatric AML cohorts stratified by **(c)** MECOM expression, **(d)** MECOM network enrichment, **(e)** MECOM NES, **(f)** clinical risk group, and **(g)** LSC17. For continuous variables in (**c)**, **(e)**, and **(g)** optimal threshold was determined by maximizing sensitivity and specificity on mortality (Youden’s J statistic). Hazard Ratios (HR) were computed from univariate cox-proportional hazard models. P values representing the result of Mantel-Cox log-rank testing are displayed. Test for trend was performed for clinical risk group stratification (>2 groups). **(h-i)** KM overall survival curves stratified by current prognostic tools and MECOM down network status. MECOM network enrichment was significantly associated with mortality independent of clinical risk group in adult (p=0.005) and pediatric (p=0.008) AML **(h)**, and independent of LSC17 score in adult (p=0.01) and pediatric (p=0.01) AML **(i)**. **(j-k)** Marginal hazard of death associated with increasing MECOM expression **(j)** and MECOM network enrichment score **(k)**, stratified by age. P-values represent significance of MECOM expression and MECOM network enrichment on survival, using multivariable cox-proportional hazards modelling, adjusted for age and sex.

Given the importance of the MECOM down gene network in HSC maintenance, we sought to determine whether expression of this network was associated with survival in AML. Using GSEA, we determined whether individual samples had enrichment or depletion of the MECOM down geneset (**Extended Data Fig. 6a-c**). Strikingly, enrichment of the MECOM down geneset was associated with worse survival in both the adult (HR: 1.52 [95%CI 1.13-2.04], pval: p=0.005) and pediatric AML cohorts (HR: 1.96 [95%CI 1.38-2.69], pval: 7.4e-5) (**Fig. 6d**).

To further characterize the effect of MECOM down gene expression on survival in AML, we generated a rank order list based on the Normalized Enrichment Score (NES) for each sample to allow for further stratification based on the degree of network enrichment. We used optimal thresholding to stratify patients based on NES and found significantly worse overall survival in patients with high MECOM NES compared to patients with low NES in both adult (HR: 1.58 [95%CI 1.18-2.11], pval: 0.0016) and pediatric (HR: 2.08 [95%CI 1.49-2.89], pval: 3.6e-5) patients (**Fig. 6e**).

Not surprisingly, stratification based on clinical risk group or LSC17 score^58^, which is enriched in leukemia stem cells and is associated with therapy resistance and poor prognosis, had significant associations with survival (**Fig. 6f,g**). Next, we sought to determine whether MECOM network enrichment identified the same subgroup of high-risk patients as clinical risk group or LSC17 score, or if it could be combined with either classification method to further stratify patient survival. We observed that 48% of adult AML and 51% of pediatric AML with adverse clinical risk features also had MECOM network enrichment. Similarly, we found that 51% of adult AML and 55% of pediatric AML with high LSC17 scores had MECOM network enrichment (**Extended Data Fig. 6d,e**). Thus, MECOM network enrichment identifies a largely unique subset of patients compared to currently available risk stratification tools.

Next, we investigated whether the addition of MECOM network enrichment to the clinical risk group or LSC17 score resulted in improved risk stratification. In the adult AML cohort, MECOM down gene set enrichment was independently associated with mortality particularly in patients with intermediate risk AML (p=0.005) (**Fig. 6h**) and high LSC17 score (p=0.01) (**Fig. 6i**). The contribution of MECOM network enrichment to clinical risk grouping was even more striking in the pediatric AML cohort in which MECOM network enrichment was significantly associated with mortality independent of clinical risk group (p=0.008) (**Fig. 6h**) and, separately, independent of LSC17 score (p=0.01) (**Fig. 6i**). These results reveal that stratification of primary AML patient samples by MECOM down network enrichment can be integrated with currently available prognostic tools to improve risk stratification for overall survival in both adult and pediatric AML. Additionally, MECOM down network enrichment was significantly associated with lower event-free survival, independent of clinical risk group and LSC17 score in pediatric AML (p=1.72e-6 and p=5.62e-5, respectively) (**Extended Data Fig. 6f-j**).

Finally, we calculated marginal hazard ratios to directly evaluate the association of MECOM expression or MECOM network NES with overall survival. We observed a modest effect of incremental increases of MECOM expression on the marginal HR of survival in adult and pediatric AML (**Fig. 6j**), and a much more significant effect of incremental increases in MECOM NES (**Fig. 6k**). Together, these data reveal that the MECOM down regulatory network is highly enriched in a subset of adult and pediatric AMLs with poor prognosis, and can be integrated with currently available prognostic tools to improve risk stratification for patients with AML.

### Validation of MECOM addiction in a subset of high-risk AMLs

Following our observations that the MECOM down gene network has prognostic significance for AML patients, we sought to further study this network in AML cell lines. First, we examined 44 AML cell lines from the Cancer Cell Line Encyclopedia (CCLE) and stratified them based on *MECOM* expression (**Extended Data Fig. 7a**). GSEA analysis of M4 AML cell lines from CCLE revealed enrichment of the MECOM down genes and depletion of the MECOM up genes in the *MECOM*-high expressing samples (**Extended Data Fig. 7b,c**). Next, we compared CRISPR dependencies of *MECOM*-high and *MECOM*-low AML cell lines from CCLE. Interestingly, we observed a striking difference in essentiality of RUNX1, consistent with our findings of potential cooperativity between RUNX1 and MECOM in regulating the HSC network genes (**Extended Data Fig. 7d**).

To validate the role of the MECOM gene network in an otherwise isogenic AML background, we performed CRISPR editing of *MECOM* in the MUTZ-3 AML cell line. MUTZ-3 cells have supraphysiologic expression of *MECOM* due to an inversion of chromosome 3 leading to juxtaposition of a *GATA2* enhancer upstream of *MECOM*^59–61^. These cells maintain a population of primitive CD34^+^ blasts in culture that can differentiate into CD14^+^ monocytes (**Fig. 7a, Extended Data Fig. 7e**). *MECOM* perturbation by CRISPR editing in MUTZ-3 cells resulted in 65% edited allele frequency (**Fig. 7b**) and significant reduction in MECOM expression level (**Fig. 7c**). *MECOM* editing of MUTZ-3 cells resulted in a loss of repopulating, primitive CD34^+^ cells and an increase of mature CD14^+^ cells by day 5 after CRISPR editing (**Fig. 7d**). Loss of progenitors after MECOM perturbation was accompanied by enrichment of edited MECOM alleles as MECOM perturbed cells underwent greater expansion (**Extended Data Fig. 7f**). Maintenance of CD34^+^ cells was restored by lentiviral *MECOM* expression, but not lentiviral expression of the shorter *EVI1* isoform (**Fig. 7e**). This observation is consistent with the data from primary HSPCs in which the full-length *MECOM* isoform was better able to rescue the loss of LT-HSCs following *MECOM* perturbation compared to *EVI1* (**Extended Data Fig. 3e**). We then sought to delineate the transcriptional changes that occur in the CD34^+^ progenitor MUTZ-3 cells after *MECOM* loss by RNA sequencing. We observed significant depletion of MECOM down genes and significant enrichment of MECOM up genes (**Fig. 7f,g** **and Extended Data Fig. 7g**) in the *MECOM*-edited MUTZ-3 samples (**Supplementary Table 6**), revealing the conservation of this gene regulatory network in both hematopoietic and leukemia stem cell populations. Finally, because of the functional interaction between MECOM and CTCF in the transcriptional control of LT-HSC quiescence, we reasoned that the loss of MUTZ-3 progenitors following *MECOM* perturbation may also be dependent on CTCF. We performed dual CRISPR editing of *MECOM* and *CTCF* (**Extended Data Fig. 7h**) and observed partial rescue of the loss of CD34^+^ progenitors induced by *MECOM* perturbation alone (**Fig. 7h**). Collectively, these data reveal that the MECOM regulatory gene network co-regulated by CTCF that is fundamental to the maintenance of LT-HSCs and that is hijacked in high-risk AML cases is indispensable for MUTZ-3 AML progenitor maintenance.

**Figure 7.**
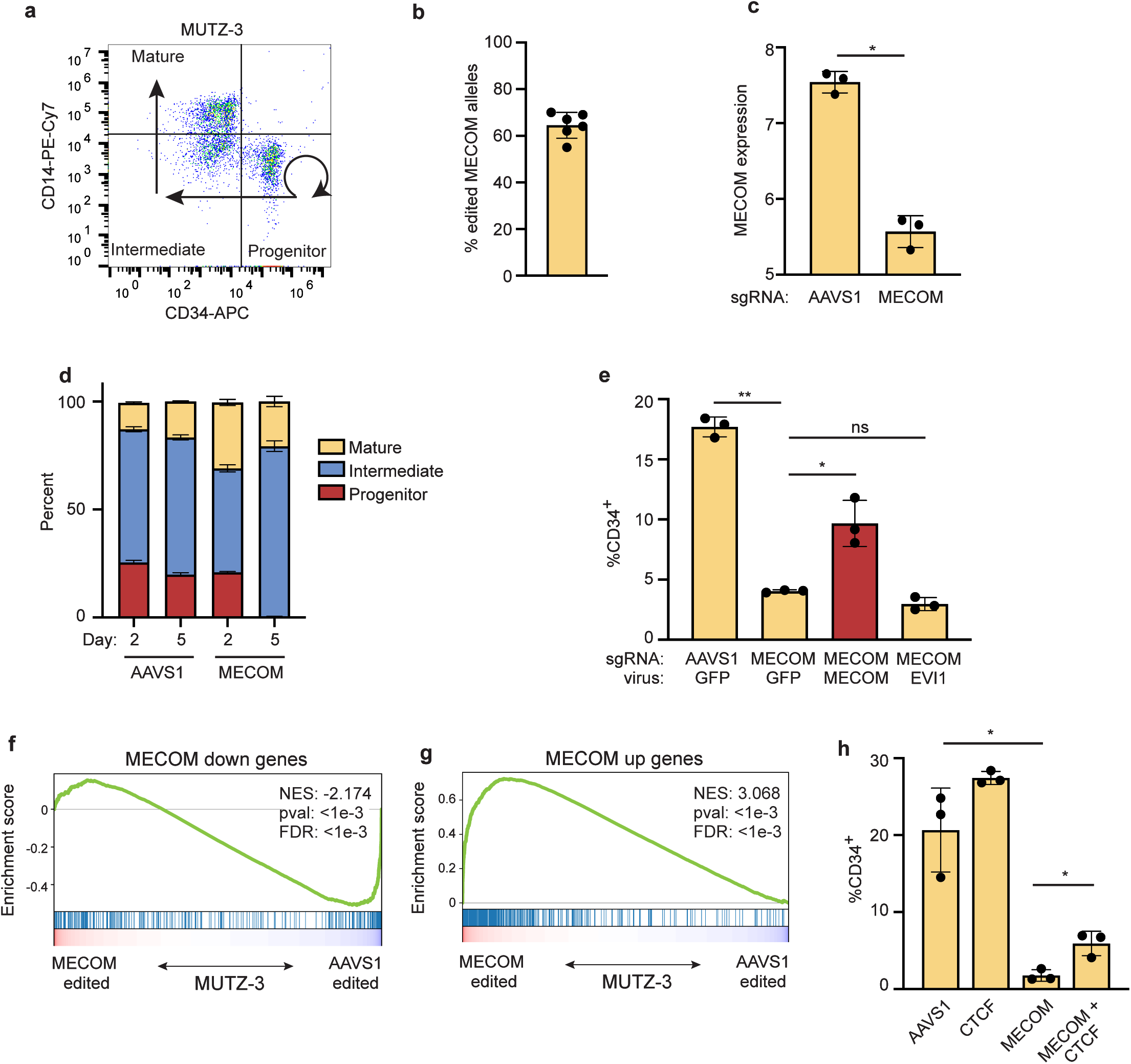
The MECOM gene regulatory network is indispensable in AML. **(a)** FACS plot showing the immunophenotype of MUTZ-3 cells. CD34^+^CD14^-^ progenitors can self-renew (curved arrow) and undergo differentiation (straight arrows) into CD34^-^CD14^-^ intermediate promonocytes and ultimately CD34^-^CD14^+^ mature monocytes. **(b)** MECOM editing in MUTZ-3 AML cells. **(c)** *MECOM* expression (log_2_ RPKM) in CD34^+^ MUTZ-3 cells. *MECOM* editing causes significant reduction in expression. *n*=3 per group. Mean is plotted and error bars show s.e.m. Two-sided Student *t*-test used. \**P* < 5e-4. **(d)** Myelomonocytic differentiation analysis of MUTZ-3 cells after CRISPR editing. Percent of cells within each subpopulation was measured by flow cytometry on days 2 and 5 after editing. *n*=3 per group. Mean is plotted and error bars show s.e.m. **(e)** Percent of MUTZ-3 cells in CD34^+^CD14^-^ progenitor population after *MECOM* editing and viral rescue as determined by flow cytometry. *n*=3 per group. Mean is plotted and error bars show s.e.m. Two-sided Student *t*-test used. ns, not significant, \**P* < 0.05, \*\**P* < 0.005. **(f-g)** GSEA of MECOM network genes in MUTZ-3 cells after *MECOM* editing. *MECOM* edited MUTZ-3 cells show enrichment of MECOM down genes **(f)**, and depletion of MECOM up genes **(g)**. **(h)** Bar graphs of the rescue of CD34^+^ by dual MECOM and CTCF perturbation. MUTZ-3 AML cells underwent CRISPR editing with the sgRNA guides depicted on the x-axis. Percent CD34^+^ cells were determined by FACS on day 4. *n*=3 per group. Mean is plotted and error bars show s.e.m. Two-sided Student *t*-test used. \**P* < 5e-2.

## DISCUSSION

Understanding the transcriptional circuitry that enables human HSCs self-renewal not only has key implications for gaining a fundamental understanding of this process, but also holds considerable promise to enable improved manipulation of such cells for therapeutic applications. For instance, with emerging advances in gene therapy and genome editing of HSCs, the ability to better maintain and manipulate these cells both *ex* and *in vivo* would be incredibly beneficial^62,63^. However, the limitations in our molecular understanding of this regulatory process have hampered such efforts and while some factors have been studied, inferences about their *in vivo* roles are often limited, particularly with the constraints of existing systems for studying and manipulating human hematopoiesis^64^.

Here, we have taken advantage of a rare experiment of nature to illuminate fundamental transcriptional circuitry that is required for human HSC maintenance *in vivo*. We have followed up on the robust human genetic observation that *MECOM* haploinsufficiency results in early onset aplastic anemia that is characterized by a paucity of HSCs. By modeling this disorder using genome editing approaches in primary hematopoietic stem and progenitor cells, we show that the functional loss of HSCs is accompanied by alterations in a network of genes critical for HSC maintenance. The identification of this gene network highlights the need to couple rigorous functional assays that can validate specific and relevant cellular vulnerabilities with integrative genomic profiling and analyses. Our results clearly demonstrate how subtle gene expression changes can translate into major deficits in HSC maintenance. Importantly, these findings are also unexpected, as there is no *a priori* reason to suspect that a network involving the regulation of hundreds of genes would be essential to maintain a stem cell population, particularly when prior functional characterization has focused on only a few key regulators^1,64^. Our findings uncover additional important regulators of HSCs that can be subject to systematic perturbational studies in the future.

Through integrative genomic analysis of this network, we have not only gained insights into the critical gene targets, but also nominated *cis*REs involved in this regulation and thereby elucidated cooperative interactions among a number of master regulator TFs involved in HSC function, including RUNX1, GATA2, and others. Moreover, we also identify an antagonistic role for CTCF in altering chromatin looping of MECOM regulatory network genes as the cells differentiate, and validate this interaction by functional and molecular rescue. Our studies illuminate the simple, yet multi-layered, molecular logic that underlies the transcriptional regulation required for human HSC maintenance.

Importantly, we have not only elucidated how a regulatory network may be altered to cause a rare genetic disorder characterized by early loss of HSCs, but we also find that this very same network is co-opted more frequently in aggressive leukemias with an extremely poor prognosis. A striking finding through our analysis is that the MECOM regulatory network serves as a better predictor of poor outcome than does *MECOM* expression itself, suggesting that some AMLs may augment MECOM function in a manner beyond expression changes. This will be an important area for future exploration. It is also notable that leukemias arising due to insertional mutagenesis following human gene therapy trials have resulted in activation of *MECOM*^65–67^. In contrast to many other insertional mutations, clones with increased *MECOM* expression often have a long latency to achieve clonal dominance, but can also result in a more aggressive disease course. Our finding that an HSC regulatory program is co-opted by increased *MECOM* expression may help explain these perplexing clinical observations. A deeper understanding of how such stem cell networks are utilized in malignant states may enable improved therapeutic approaches, while also providing opportunities to expand and manipulate non-malignant HSCs for therapeutic benefit.

## ACKNOWLEDGEMENTS

We are grateful to members of the Sankaran laboratory and numerous colleagues for valuable comments and suggestions. This work was supported by the New York Stem Cell Foundation (V.G.S.), a gift from the Lodish Family to Boston Children’s Hospital (V.G.S.), the Klarman Cell Observatory (A.R.), and National Institutes of Health Grants R01 DK103794 and R01 HL146500 (V.G.S.). R.A.V. and L.W. received support from National Institutes of Health Grant T32 HL007574. S.K.N. is a Scholar of the American Society of Hematology. V.G.S. is a New York Stem Cell-Robertson Investigator.

## AUTHOR CONTRIBUTIONS

R.A.V., L.T., F.Y., and V.G.S. conceived and designed the experiments and wrote the manuscript with input from all authors. R.A.V., L.T., L.D.C., B.C., C.F., S.K.N., L.W., X.L., and K.T. performed functional studies and provided interpretation. F.Y. and L.T. performed the computational analyses. F.Y. designed and developed HemeMap. A.R. and V.G.S. provided supervision and overall project oversight.

## EXTENDED DATA FIGURE LEGENDS

**Extended Data Figure 1. Modeling *MECOM* haploinsufficiency in human CD34^+^ HSPCs.**

**(a)** Schematic of the *MECOM* locus annotated with the location of sgRNAs (sg1-sg9) tested for efficiency of *MECOM* editing. The binding site of sg8 (underlined) which is used in subsequent studies, and clinical mutations described in *MECOM* haploinsufficient bone marrow failure (red) are indicated.

**(b)** Predicted partial protein structure of the MECOM zinc finger domain with mutated residues shown as spheres. These mutations are expected to disrupt the structure of the zinc finger, either through abrogation of Zn coordination (H751, C766) or tethering between the ZnF (R750, R778).

**(c)** Percent modified alleles after transient transfection of sgRNA and Cas9 plasmids into 293T cells. Editing frequency was detected at 72 hours after transfection by Sanger sequencing and ICE analysis. Mean is plotted and error bars show s.e.m.

**(d)** Comparison of Sanger sequencing followed by ICE analysis and Next Generation Sequencing (NGS) for the detection of CRISPR edits. *AAVS1* (blue) and *MECOM* (red) edited samples were analyzed by ICE and NGS in parallel.

**(e)** *MECOM* editing in human CD34^+^ HSPCs after RNP delivery by nucleofection. Editing frequency was detected at 48 hours by Sanger sequencing of genomic DNA. Transcription of edited *MECOM* alleles was determined by qRT-PCR from bulk RNA of HSPCs at 48 hours. Mean is plotted and error bars show s.e.m.

**(f)** *MECOM* expression following CRISPR editing. *MECOM* expression (normalized to *GAPDH*) in bulk HSPCs was detected by qRT-PCR (*n*=3 per time point; three biologically independent experiments) and was normalized to expression in the AAVS1 edited sample on the same day. Mean is plotted and error bars show s.e.m. Two-sided Student *t*-test used. \**P*< 1e-3.

**(g)** *MECOM* expression in LT-HSCs. *MECOM* expression (normalized to *GAPDH*) was detected by qRT-PCR (*n*=3 per group; three biologically independent experiments) in bulk CD34^+^ HSPCs and in LT-HSCs sorted on day 3 after CRISPR editing. Mean is plotted and error bars show s.e.m. Two-sided Student *t*-test used. \**P* < 0.01.

**(h)** Expansion of LT-HSCs in culture. HSPCs were cultured in the presence (*n*=2) or absence (*n*=2) of the HSC self-renewal agonist UM171. Percent of LT-HSCs was determined by FACS as in **Fig. 1e** and was used to calculate the total LT-HSC number. Cells were supplemented with fresh media every 2 days.

**(i)** Expansion time course of bulk CD34^+^ HSPCs following CRISPR editing. HSPCs were thawed into HSC media containing 35nM UM171 and underwent CRISPR editing 24 hours later. Cells were counted daily by trypan blue exclusion starting on day 2 after CRISPR editing and media was added to maintain equal confluency. *n*=3 per group. Mean is plotted and error bars show s.e.m. Two-sided Student *t*-test used. \**P* < 5e-3.

**(j)** Stacked bar graph of cell cycle status of bulk HSPCs and HSC (HSC: CD34^+^CD45RA^-^ CD90^+^CD133^+^) as determined by Edu incorporation and 7-AAD staining. On day 5 after CRISPR editing, cells were incubated with Edu for 2 hours, then fixed and permeabilized prior to 7-AAD and cell surface staining. Comparing *AAVS1*-edited (A) and *MECOM*-edited (M) samples, there was no difference in the proportion of cells in G0/G1 (Edu^-^/2n DNA content), S (Edu^+^), or M (Edu^-^/>2n DNA content) in bulk CD34^+^ cells or CD34^+^CD45RA^-^CD90^+^ HSCs. *n*=3 per group.

**(k)** Stacked bar graph of cell cycle status of LT-HSCs as determined by transcriptional signatures of single-cell LT-HSCs. UCB CD34^+^ underwent CRISPR perturbation of *MECOM* or *AAVS1* and were maintained in HSC media. On day 4 after editing, LT-HSCs were sorted and 10x scRNA sequencing was performed. There was no difference in cell cycle state in LT-HSCs following *AAVS1* or *MECOM* editing.

**(l)** Analysis of cell expansion following CRISPR editing. *AAVS1* or *MECOM* edited HSPCs were labeled with CFSE and successive generations of cell divisions were determined by CFSE signal intensity on day 5 which was used to calculate the replication index. Mean of three independent experiments is plotted and error bars show s.e.m. Two-sided Student *t*-test used. \**P* < 5e-2.

**(m)** Stacked bar plots of colony forming assay comparing *MECOM* edited adult CD34^+^ HSPCs (*n*=6) to *AAVS1* edited controls (*n*=3). CFU-GEMM, colony-forming unit (CFU) granulocyte erythroid macrophage megakaryocyte; CFU-GM, CFU granulocyte macrophage; CFU-M, CFU macrophage; CFU-G, CFU granulocyte. Mean colony number is plotted and error bars show s.e.m.

**(n-o)** NGS of *MECOM* in human HSPCs following CRISPR editing, prior to xenotransplantation **(n)**, and after harvest from bone marrow at 16 weeks of one representative mouse **(o)**. Sequences present at frequencies >0.5% are displayed.

**(p)** Analysis of bone marrow of mice at week 16 following transplantation of *MECOM*-edited (*n*=5) and *AAVS1*-edited (*n*=3) adult HSPCs. Mean is indicated by black line and each data point represents one mouse. Two-sided Student *t*-test used. \**P* < 0.05.

**(q)** Analysis of the *MECOM* locus of human cells harvested from mice following primary or secondary xenotransplantation. Half of the primary recipient mice (4/8) had human chimerism >0.25% (circles) and the other half had chimerism <0.25% (triangles) but had human *MECOM* sequences that were detectable by PCR. All of the secondary recipients had human chimerism <0.25% but had human *MECOM* sequences that were detectable by PCR.

**Extended Data Figure 2. Single cell RNA sequencing of LT-HSCs after MECOM editing**.

**(a-c)** UMAP plots of the normalized expression of *CD34* **(a)**, *HLF* **(b)**, and *CRHBP* **(c)** in phenotypic LT-HSCs. The combined expression of these three genes defines the HSC signature in **Fig. 2a,b**.

**(d-e)** Louvain clustering of LT-HSCs **(d)**; Bar graph of the ratio of cells in Louvain cluster 1 and 2 following *AAVS1* or *MECOM* editing **(e)**.

**(f)** Volcano plot projection of the data from **Fig. 2e,f** displaying the small but significant fold changes in gene expression of MECOM down genes (log_2_ fold change < -0.05) and MECOM up genes (log_2_ fold change > 0.05) with *p*-value <1e-20. Log_2_fold change of MPO expression is out of scale of the axis and is noted by a red arrow.

**(g-h)** Box plots showing expression of a subset of MECOM down **(g)** and MECOM up **(h)** genes in a representative random permutation of cohort assignments, demonstrating no difference in gene expression. Gray dots show imputed gene expression in single cells.

**(i)** Scatter plot of gene expression in LT-HSCs enriched for the transcriptional HSC signature compared to bulk LT-HSCs. Expression differences between *MECOM* and *AAVS1* edited LT-HSCs were calculated and MECOM down and MECOM up genes are plotted. Correlation was calculated using Spearman’s rank correlation test.

**Extended Data Figure 3. Lentiviral expression of *MECOM* rescues LT-HSCs but does not reverse upregulation of MECOM up genes.**

**(a)** Schematic of lentiviral vector for increased *MECOM* expression. MECOM sgRNA binding site is shown in bold, and wobble mutations introduced by PCR are indicated. LTR, long terminal repeat; IRES, internal ribosome entry site.

**(b)** Edited allele frequency of intended endogenous *MECOM* locus and *MECOM* cDNA after viral integration. Editing and infection were performed as in **Fig. 3a**. Integrated viral cDNA was amplified using a forward primer in the cDNA sequence and reverse primer in the IRES sequence. Mean is plotted and error bars show s.e.m.

**(c)** FACS plots for LT-HSC detection after *MECOM* editing and rescue. Gating strategy as in **Fig. 1e**. Percentages show the mean (± s.e.m) of three independent experiments. GFP ratio (**Fig. 3e**) is defined as the ratio of GFP^+^ cells in LT-HSC population (column 4) to GFP^+^ cells in the bulk population (column 5).

**(d)** Cell expansion after *MECOM* editing and rescue. Increased expansion of HSPCs after *MECOM* editing is not reversed by viral *MECOM* expression. AAVS1, edited at *AAVS1*, infected with GFP virus; MECOM, edited at *MECOM*, infected with GFP virus; rescue, edited at *MECOM*, infected with *MECOM* virus, *n*=3 for each group. Mean is plotted and error bars show s.e.m. Two-sided Student *t*-test used. \**P* < 5e-3.

**(e)** Bar graph of the effect of MECOM isoform overexpression on the maintenance of LT-HSCs. HSPCs were edited at *AAVS1* (yellow) or *MECOM* (red) and infected with lentivirus encoding GFP or MECOM isoforms as displayed. The percentage of LT-HSCs was determined by FACS. Mean is plotted and error bars show s.e.m.

**(f-g)** GSEA of MECOM up genes after editing and rescue in bulk LT-HSCs. **(f)** MECOM up genes are more highly enriched in AAVS1 samples in bulk in contrast to data from single cell analysis (**Fig. 2f**). **(g)** MECOM up genes are further increased after MECOM viral infection.

**Extended Data Figure 4. Establishment of a *cis*-regulatory network in HSCs.**

**(a)** Schematic view demonstrating different types of functional interactions between *cis*-regulatory elements and genes. HemeMap predicts these interactions by integration of multiomics data including RNAseq, ATACseq and promoter capture-HiC (PC-HiC) data across 16 or 18 hematopoietic cell types.

**(b)** Bar graph showing the overlap between genomic interactions nominated by HemeMap and experimentally-defined interactions. More than half of the direct interactions nominated by PC-HiC and RNA-ATAC correlations were supported by evidence from Hi-C interactions in HSPCs.

**(c)** Correlation of *cis*RE-gene interaction strength with gene expression in HSCs. HemeMap scores were calculated for each *cis*REs-gene interaction and HemeMap interactions were arranged by increasing scores and grouped evenly into 50 bins. Median gene expression in each bin is depicted (bars). The median expression of a randomly sampled equal-sized gene set is shown (dots).

**(d)** Distribution of HemeMap scores in HSCs. To construct the HSC-specific regulatory network, significant interaction scores >8.91 were included. Significance threshold was determined by Chi-square distribution.

**(e)** Comparison of interaction strengths. *cis*REs containing ETS footprint were significantly associated with stronger HemeMap scores than those without. P-values as shown were calculated using the Wilcoxson signed-rank test.

**(f-g)** Analysis of TF footprint co-occurrence in the *cis*REs associated with MECOM down genes **(f)** and MECOM up genes **(g)**, respectively. The frequency of occurrence and *P* values were calculated using a hypergeometric test. The color and size of dots are proportional to statistical significance.

**Extended Data Figure 5. CTCF-mediated looping of MECOM down genes in HSCs.**

**(a)** Boxplots depict the quantitative difference of CTCF ChIP-seq signals between CD34^+^ HSPCs and lineage-committed cells from **Fig. 5d**. The normalized signals of CTCF ChIP-seq signals of 50 bp regions centered on CTCF footprints were calculated and compared. The significance was determined using Wilcoxson signed-rank test, *** *P*<5e-6.

**(b-d)** Boxplots displaying the chromatin accessibility of CTCF-associated c*is*REs during hematopoietic differentiation. MECOM down *cis*REs that contain a CTCF footprint are associated with progressively less chromatin accessibility during differentiation along the **(b)** erythroid, **(c)** myeloid, and **(d)** lymphoid lineages.

**(e)** Chromatin interactions of MECOM down genes based on the presence and orientation of CTCF footprint. 448 chromatin interactions involving MECOM down genes were identified and were categorized as: (1) no CTCF footprint detected at either anchor (2) CTCF present both anchors in same orientation (3) CTCF present both anchors in opposite orientation (4) CTCF present at only one anchor.

**(f)** Bar graphs of CRISPR editing frequencies in human HSPCs. Cells that underwent dual CRISPR perturbation of *MECOM* and *CTCF* had editing similar frequencies compared to single-edited cells. *n*=3 per group. Mean is plotted and error bars show s.e.m.

**(g)** Bar graphs of total cell number following CRISPR editing. Increased expansion of HSPCs following MECOM perturbation was seen as in **Extended Data Fig. 1i** and was rescued by dual *MECOM* and *CTCF* perturbation. *n*=3 per group. Mean is plotted and error bars show s.e.m. Two-sided Student *t*-test used.* *P*<5e-2.

**(h-i)** GSEA of MECOM down genes **(h)** and MECOM up genes **(i)** in bulk LT-HSCs after *MECOM* perturbation compared to *AAVS1* perturbation. MECOM down genes are depleted and MECOM up genes are enriched following *MECOM* editing.

(**j-k**) Expression of MECOM down genes that are associated with CTCF loops **(j)** and those not associated with CTCF loops **(k)**, following either *MECOM* perturbation alone or dual *MECOM* and *CTCF* perturbation. P-values as shown were calculated using the Wilcoxson signed-rank test.

**Extended Data Figure 6. MECOM down gene network enrichment is independently associated with overall and event-free survival**

**(a-c)** GSEA of MECOM down genes in primary AML patient samples from TCGA. For each patient sample, expression of every gene was compared to its average expression from all TCGA patient samples, and GSEA was performed to assess for enrichment of MECOM down genes. Representative plots of three individual patients are shown. **(a)** Patient 2896 had enrichment of MECOM down genes and an overall survival of 230 days. **(b)** Patient 3011 had depletion of MECOM down genes and an overall survival of 2450 days. **(c)** Patient 2982 had no significant enrichment or depletion of MECOM down genes and an overall survival of 1110 days.

**(d-e)** Stacked bar graph showing proportion of patients with MECOM network enrichment or depletion following stratification by clinical risk group or LSC17 score in adult **(d)** or pediatric AML **(e)**.

**(f-j)** KM event-free survival curves for the pediatric AML cohort stratified by **(f)** MECOM expression, **(g)** MECOM network enrichment, **(h)** MECOM NES, **(i)** clinical risk group, and **(j)** LSC17. For continuous variables in **(f)**, **(h)**, and **(j)** the optimal threshold was determined by maximizing sensitivity and specificity on mortality (Youden’s J statistic). Hazard Ratios (HR) were computed from univariate cox-proportional hazard models. P values representing the result of Mantel-Cox log-rank testing are displayed. Test for trend was performed for clinical risk group stratification (>2 groups).

**Extended Data Figure 7. Evaluation of the MECOM gene network in high-risk AML.**

**(a)** Violin plots showing *MECOM* expression in AML samples from CCLE. AML samples were stratified by *MECOM* expression (log_2_ RPKM +1). Low, <1 (*n*=31); High≥1 (*n*=13). Mean is plotted and dashed lines indicate quartiles.

**(b-c)** GSEA of MECOM network genes in CCLE M4 AML samples. MECOM high AMLs show enrichment of MECOM down genes **(b)**, and depletion of MECOM up **(c)** genes compared to MECOM low AMLs.

(**d)** Volcano plot showing differential CRISPR dependencies of CCLE AMLs stratified by *MECOM* expression. Average CRISPR dependencies for the CCLE AML cohorts as defined in **Extended Data Fig. 7a** were determined using CERES and effect size was calculated by comparing dependency scores of *MECOM* high and *MECOM* low AMLs. Effect size of 0 indicates no difference in essentiality whereas negative effect size indicates higher essentially in *MECOM* high AML.

(**e)** FACS plots showing the differentiation of MUTZ-3 cells after CD34 selection. CD34+ MUTZ-3 cells were magnetically separated using the EasySep Human CD34 Positive Selection Kit II, cultured in MUTZ-3 media, and analyzed by flow cytometry at the indicated timepoints.

(**f)** Time course of edited allele frequency in MUTZ-3 AML. Genotyping was performed in bulk MUTZ-3 cells following CRISPR editing at AAVS1 (blue) or MECOM (red). Mean is plotted and error bars show s.e.m. Missing error bars are obscured by the icons.

(**g)** Violin plot of differential gene expression in CD34^+^ MUTZ-3 cells following *MECOM* perturbation. MECOM down genes are significantly depleted and MECOM up genes are significantly enriched in *MECOM* edited samples compared to AAVS1 edited samples, unlike a set of randomly selected genes. Two-sided Student *t*-test used. **** *P* < 1e-4.

(**h)** Bar graphs of CRISPR editing frequencies in MUTZ-3 AML. Cells that underwent dual CRISPR perturbation of *MECOM* and *CTCF* had similar editing frequencies compared to single-edited cells. Mean is plotted and error bars show s.e.m.

## SUPPLEMENTARY TABLES

**Supplementary Table 1. *MECOM* mutations in bone marrow failure.** Summary of genetic and clinical data of patients with *MECOM* haploinsufficiency described in the literature.

**Supplementary Table 2. MECOM network genes.** Differentially expressed genes after MECOM editing as determined by MAST. Pct.1 and pct.2 indicate the percentage of cells expressing the gene in MECOM or AAVS1 edited samples, respectively.

**Supplementary Table 3. Rescue of MECOM network genes in LT-HSCs.** Normalized expression of MECOM down and MECOM up genes in LT-HSCs analyzed in bulk. *n*=3 for each group.

**Supplementary Table 4. HemeMap interactions of MECOM down genes.**

**Supplementary Table 5. HemeMap interactions of MECOM up genes.**

**Supplementary Table 6. MECOM regulated gene network in MUTZ3 AML.** Normalized expression of MECOM down and MECOM up genes in MUTZ-3 cells after editing. *n*=3 for each group.

## METHODS

### Cell line and primary cell culture

HSPCs were purified from discarded umbilical cord blood samples of healthy male or female newborns using the EasySep Human CD34 Positive Selection Kit II following pre-enrichment using the RosetteSep Pre-enrichment cocktail (Stem Cell Technologies) and mononuclear cell isolation on Ficoll-Paque (GE Healthcare) density gradient. Cells were cryopreserved for later use. G-CSF mobilized adult CD34^+^ HSPCs and were purchased (Fred Hutchinson Cancer Research Center). Thawed cells were cultured at 37°C and 5%O_2_ in serum-free HSC media comprised of StemSpan II medium (Stem Cell Technologies) supplemented with CC100 cytokine cocktail (Stem Cell Technologies), 100ng/ml TPO (Peprotech) and 35nM UM171 (Stem Cell Technologies). Confluency was maintained between 2e5-1e6 cells/ml.

MUTZ-3 cells (DSMZ) were cultured at 37°C in alpha-MEM (Life Technologies) supplemented with 20% FBS, 20% conditioned media from 5637 cells (ATCC)^68^ and 1% penicillin/streptomycin. Confluency was maintained between 7e5-1.5e6/ml.

293T cells were cultured at 37°C in DMEM (Life Technologies) supplemented with 10% FBS and 1% penicillin/streptomycin.

### Mouse model

NOD.Cg-*Kit*^W-41J^*Tyr*^+^*Prkdc*^scid^*Il2rg*^tm1Wjl^(NBSGW) mice were obtained from Jackson Laboratory (Stock 026622)^34^. Littermates of the same sex were randomly assigned to experimental groups. NBSGW were interbred to maintain a colony of animals homozygous or hemizygous for all mutations of interest. All animal experiments were approved by the Boston Children’s Hospital Institutional Animal Care and Use Committee.

### CRISPR editing and analysis

Electroporation was performed on day 1 after thawing HSPCs using the Lonza 4D Nucleofector with 20 µl Nucleocuvette strips as described^36,69^. Briefly, ribonucleoprotein (RNP) complex was made by combining 100pmol Cas9 (IDT) and 100pmol modified sgRNA (Synthego) targeting MECOM (CAAGGTCTGCAAACCTAACA), AAVS1 (GGGGCCACTAGGGACAGGAT) or CTCF (CAATTCTCCACTGGTCACAA) and incubating at room temperature for 15 minutes. 2e5-4e5 HSPCs resuspended in 20 µl P3 solution were mixed with RNP and underwent nucleofection with program DZ-100. For samples that underwent dual perturbation, total amounts of 100pmol Cas9 and 100mol sgRNA (50 pmol each guide) were used. Cells were returned to HSC media and editing efficiency was measured by PCR at 48 hours after electroporation, unless otherwise indicated. First, genomic DNA was extracted using the DNeasy kit (Qiagen) or both DNA and RNA were extracted using the AllPrep DNA/RNA Mini kit (Qiagen) according to the manufacturer’s instructions. Genomic PCR was performed using Platinum II Hotstart Mastermix (Thermo) and edited allele frequency was detected either by Sanger sequencing and analyzed by ICE^70^, or NGS and analyzed with Crispresso2^71^. The following primer pairs were used: MECOM-ICE (forward: ACATCAACCCAGAATCAGAAAC; reverse: GGAAAAGGAAGGCTGCAAAG), MECOM-NGS (forward: AGAAATGTGAGTTCCATGCAAGA; reverse: AGCAAATATCATTGTCAGACCTGT). CTCF (forward: CAGCGGATTCAGATGGGTAA; reverse: TCACCGTTTTAGCCAGGATG). The effect on MECOM mRNA after editing was detected by qRT-PCR using SYBR green (Biorad) after cDNA synthesis with iScript (Biorad).

MUTZ-3 cells were edited as above with the following modification: cells were resuspended in 20 µl SF solution and program EO-100 was used for electroporation.

### Viral constructs and transduction

*MDS* and *EVI1* cDNA were synthesized from mRNA of human HSPCs using the following primers: MDS (forward: CGTACTCGAGGCCGCCACCATGAGATCCAAAGGCAGGGCAA; reverse: TACGGAATTCTCACTCCCATCCATAACTGGGGTCT), EVI1 (forward: CGTACTCG AGGCCGCCACCATGATCTTAGACGAATTTTACAATG; reverse: TACGGAATTCTCATACGTGGCTTATGGACTGG). *MECOM* cDNA was synthesized using MDS-F and EVI1-R primers. Wobble mutations were introduced to disrupt the sgRNA binding site using the following primers EVI1-F and wobble reverse (GTGCCGAGTGAGATTCGCGGATCTAGGAAAAAT) and wobble forward (ATTTTTCCTAGATCCGCGAATCTCACTCGGCAC) with EVI1-R, followed by overlap PCR of the two fragments. Primers included restriction enzyme sites to allow for cloning using EcoRI and XhoI into the HMD IRES-GFP backbone^72^.

To produce lentivirus, approximately 24 hours prior to transfection, 293T cells were seeded in 10cm plates. Cells were co-transfected with 10µg pΔ8.9, 1µg VSVG, and 10µg HMD vector variant using calcium phosphate. Media was changed the following day and viral supernatant was harvested at 48 hours post-transfection, filtered with a 0.45um filter and concentrated by ultracentrifugation at 24,000 r.p.m. for 2 hours at 4°C.

For lentiviral rescue experiments, 24 hours after CRISPR nucleofection, 1e5 HSPCs were transduced at a multiplicity of infection (MOI) of 10, with HMD empty, MDS, EVI1 or MECOM virus in 12 well plates with 8µg/ml of polybrene (Millipore), spun at 2,000 r.p.m. for 1.5 hours at room temperature and incubated in the viral supernatant overnight at 37°C. Virus was washed off 16 hours after infection.

MUTZ-3 cells were transduced at an MOI of 1 by spinfection at 2,500 r.p.m. for 1.5 hours at room temperature and were incubated in the viral supernatant overnight. Virus was washed off 16 hours after infection. MUTZ-3 cells underwent viral transduction first, followed by CRISPR editing at 48 hours post-infection.

### Transplantation assays

Non-irradiated NBSGW mice (between 4–8 weeks of age) were tail vein injected with UCB or adult CD34^+^ HSPCs (1-2e5 cells) on day 3 after CRISPR editing. Peripheral blood was sampled monthly by retro-orbital sampling and animals were sacrificed at 16 weeks for bone marrow evaluation. Bone marrow cells were collected by flushing of both femurs and tibias. Secondary transplantations were performed by directly transplanting 60% of total BM cells from primary recipients into secondary non-irradiated NBSGW recipients. Human chimerism was assessed by evaluation of the bone marrows of secondary recipients at 16 weeks by flow cytometry as below and MECOM sequencing was performed as above.

### Flow cytometry and cell sorting

Cells were washed with PBS and stained with the following panel of antibodies to quantify and enrich for LT-HSCs: anti-CD34-PerCP-Cy5.5 (Biolegend, 343612), anti-CD45RA-APC-H7 (BD, 560674), anti-CD90-PECy7 (BD, 561558), anti-CD133-super bright 436 (Ebioscience, 62-1338-42), anti-EPCR-PE (Biolegend, 351904) and anti-ITGA3-APC (Biolegend, 343808). LT-HSCs were defined by the following immunophenotype: CD34^+^CD45RA^-^CD90^+^ CD133^+^ITGA3^+^EPCR^+^^28^. Three microliters of each antibody were used per 1e5 cells in 100µl. Total LT-HSC numbers were calculated as a product of the frequency of LT-HSCs by flow cytometry and total cell number in culture.

Human cell chimerism after xenotransplantation was determined by staining with anti-mouse CD45-FITC (Biolegend, 103108) and anti-human CD45-APC (Biolegend, 368512). Human cell subpopulations were detected in the bone marrow of transplanted mice using the following antibodies: anti-human CD45-APC (Biolegend, 368512), anti-human CD3-Pacific Blue (Biolegend, 344823), anti-human CD19-PECy7 (Biolegend, 302215), anti-human CD11b-FITC (Biolegend, 301330), anti-human CD41a-FITC (Ebioscience, 11-0419-42), anti-human CD34-Alexa 488 (Biolegend, 343518) and anti-human CD235a-APC (Ebioscience, 17-9987-42). Aliquots were stained individually for CD34 and CD235, or with CD45 in conjunction with the other lineage-defining markers. Mice with human cell chimerism less than 2% in the bone marrow were excluded from subpopulation analysis.

MUTZ3 cells were stained with anti-CD34-APC (Biolegend, 343607) and anti-CD14-PECy7 (Biolegend, 367112).

Flow cytometric analyses were conducted on Becton Dickinson (BD) LSRII, LSR Fortessa or Accuri C6 instruments and all data were analyzed using FlowJo software (v.10.6). Fluorescence-activated cell sorting (FACS) was performed on BD Aria and samples were collected in PBS containing 2% BSA and 0.01% Tween for immediate processing for sequencing on the 10x Genomics platform. Alternatively, single cells were sorted into PCR plates containing 5 µl Buffer RLT Plus (Qiagen) with 1% BME and immediately frozen at -80°C for G&T sequencing.

### Cell cycle analysis

For cell cycle analyses, on day 5 after CRISPR editing, cells were incubated with Edu (Thermo, C10634) for 2 hours, then fixed and permeabilized prior to cell surface staining as per the manufacturer’s recommendations. Multipotent progenitors were defined by the following immunophenotype: CD34^+^CD45RA^-^CD90^+^ CD133^+^. Pegasus 1.0 (https://github.com/klarman-cell-observatory/pegasus) in the Terra environment (https://app.terra.bio/#) was used to determine the expression of transcriptional signatures of cell cycle status of single LT-HSCs^92^.

Analysis of cell division was performed by CFSE labeling (Thermo, C34554). 24 hours after CRISPR editing, cells were incubated with CFSE, washed and subjected to flow cytometric analysis to establish a baseline. Five days later, cells were again analyzed by flow cytometry and the number of cells in each divisional generation was determined by proliferation modeling in Flowjo v10.8.0. Replication index is a measure of the expansion of the cells that have undergone at least one cell division.

### Colony forming unit cell assays

Three days after RNP electroporation, 500 CD34^+^ HSPCs were plated in 1 ml methylcellulose media (H4034, Stem Cell Technologies) in triplicate unless otherwise noted. Primary colonies were counted after 14 days.

### 10x single-cell RNA sequencing

A suspension of 11,000 *AAVS1*-edited LT-HSCs and a suspension of 16,000 *MECOM*-edited LT-HSCs were loaded into two lanes of 10x RNA 3’ V3 kit (10x Genomics) according to the manufacturer’s guidelines. Two libraries were constructed with distinct i7 barcodes, pooled in equal molecular concentrations and sequenced on one lane of Hiseq (Illumina) according to the manufacturer’s protocol.

### Bulk RNA sequencing

Total RNA was extracted using the Qiagen RNeasy Micro Kit (Cat. No: 74004) or using the 2.2x RNAClean XP kit (Beckman A63987) from ∼1000 cells sorted in 25 µl Buffer RLT Plus with 1% BME. Then we proceeded with the Smart-Seq2 protocol from the RT step using 10 ng of RNA^73^. The whole transcriptome amplification step was set at 10 cycles. 15 bulk RNA libraries were pooled at equal molecular concentration and sequenced using the NextSeq 550 High Output kit (Illumina) with 35 paired-end reads.

### Genome & transcriptome sequencing

Plates of sorted LT-HSCs were thawed from -80°C on ice, and an equal volume of prepared 2x Dynabeads was added. Samples were incubated at 72°C for 1 min, then 56°C for 2 min, followed by 10 min at 25°C, to allow for mRNA hybridization. Plates were placed on a magnet for 2 min and 8 µl of the supernatant containing genomic DNA (gDNA) was transferred into a new plate. Beads were washed twice in 10 µl of cold 1X Hybridization Buffer and once in PBS + RNase Inhibitor. All washes were transferred to the gDNA plate. Once PBS was removed, Dynabeads were immediately resuspended in 7.34 µl of SmartSeq2 Mix 1, and the plate was incubated at 80°C for 3 min. The plate was immediately placed on the magnet and the supernatant containing mRNA was rapidly transferred into a new plate on ice. 2.66 µl of SmartSeq2 Mix 2 was added. At this point, we proceeded with the Smart-Seq2 protocol from the RT step^73^. The whole transcriptome amplification step was set at 23 cycles. gDNA which was present in the pooled supernatant/wash buffer was precipitated on DNA SPRI beads at a 0.6X ratio, and eluted in 10 µl MDA Hyb buffer, denatured at 95°C for 3 min, and cooled on ice. Then 5 µl of Phi29 Mix was added, and the mix was incubated at 45°C for 8 hours. The reaction was deactivated at 65°C for 5 min. The MDA plate was stored at -20°C. 8 plates of mRNA libraries were sequenced using the Nextseq550 high output kit (Illumina) with 35 paired-end reads according to the manufacturer’s recommendations. To genotype each cell based on MECOM editing status, MECOM from gDNA and WTA was amplified by PCR, and libraries were constructed, pooled and sequenced using the Miseq 300 cycle kit (Illumina) according to manufacturer’s protocol with 150 paired-end reads.

## QUANTIFICATION AND STATISTICAL ANALYSIS

### Protein structure prediction

The MECOM sequence corresponding to amino acids 700-900 was submitted to the I-TASSER server for homology modeling^74^. The predicted structure of the zinc finger domain was rendered and visualized using PyMOL.

### Bulk RNA data analysis

Fastq files demultiplexed by bcl2fastq from bulk RNA-seq run were uploaded to Terra and processed with the Cumulus pipeline for bulk RNA-seq^75^ to get gene counts and gene isoform matrices. Human reference genome GRCh38 and gene annotation reference Homo_sapiens.GRCh38.93.gtf were used in all the RNA analysis.

### Single cell RNA data analysis

BCL files generated by scRNA-seq were uploaded to Terra and processed with the Cumulus pipeline for 10x single cell RNA data and SmartSeq2^75^ to get gene matrices. Human reference genome GRCh38 and gene annotation reference Homo_sapiens.GRCh38.93.gtf were used in all the RNA analysis. For 10x data, doublets were filtered out, and cells that contained reads for 500 to 8000 genes with the percent of mitochondrial genes <20% were included in the analysis; cells were not filtered based on UMI counts. For SmartSeq2 data, Scanpy^76^ was used to integrate all plates, and perform batch correction and normalization. Cells that contained reads for 2,000 to 20,000 genes with the percent of mitochondrial genes <20% were included. Genes expressed in at least 0.05% of cells were included. Scanorama^77^ was used for batch correction. SmartSeq2 and 10x data were integrated and batch correction was performed on donor, technology, and process batch with a Python version of Harmony^78^.

### MECOM genotyping in G&T data

*MECOM* editing was determined by CRISPResso2^71,75^. Genotyping from gDNA and from cDNA was combined for the same cell, and cells that contained both an edited allele and a wildtype allele were defined as heterozygous. Genotyping annotation was integrated into gene matrix meta data.

### Differential expression analysis

DE analysis was done by Seurat 4.0 with the function FindMarkers pipeline in the 10x single cell RNA data to compare *AAVS1*- and *MECOM*-edited LT-HSCs. The fold change threshold for significant gene expression was 0.05 on log_2_ scale, ident.1 was *AAVS1*-edited cells, ident.2 was *MECOM*-edited cells, and the test algorithm was MAST. Permutation analysis was performed by randomly assigning single cells to one of two groups irrespective of the initial experimental group and repeating DE analysis. 100 independent permutations were performed.

### Pseudo bulk analysis

Raw counts from single LT-HSCs that passed the quality control from each experimental condition (*AAVS1* or *MECOM* edited) were aggregated to generate pseudo bulk data for each group. Genes that did not reach the detection ratio cutoff used in the single-cell differential gene expression discovery were removed from the pseudo-bulk analysis. Log_2_ fold change between groups was calculated and correlation with gene expression data from single cells was calculated by Spearman’s rank correlation.

### HSC signatures in Immune Cell Atlas

Pegasus 1.0 was used to determine the expression of the HSC signature (CD34, HLF, CRHBP)^36^ in umbilical cord samples from the Immune Cell Atlas (https://data.humancellatlas.org/explore/projects/cc95ff89-2e68-4a08-a234-480eca21ce79).

### Gene signature enrichment during hematopoiesis

We measured the enrichment of the MECOM down or MECOM up genesets during hematopoiesis, using bulk RNA-seq datasets across 20 hematopoietic sub-populations^39^. The observed expression *y_i,j_* for the tested gene set *i* in cell type *j* was calculated by taking the mean expression of genes in the list. We performed 1,000 permutations in which we sampled gene sets with the same number of genes as the tested geneset. The expected expression 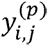, for permuted geneset *i* in cell type *j* was calculated by taking the mean expression of genes in the list. The enrichment *z* for geneset *i* in cell type *j* was computed as follows:

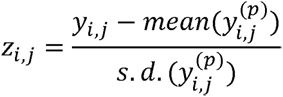

where the mean and variance of 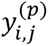 are taken over all values of *p*(*p* ɛ (1,2,…, 1000).

### Gene set enrichment analysis

We used GSEApy (https://github.com/zqfang/GSEApy) for all GSEA analyses to determine the enrichment of MECOM network genes following MECOM editing and rescue, and in the TCGA and CCLE datasets that were stratified based on MECOM expression or overall survival. Significant enrichment of the geneset was determined using *t*-test for MECOM rescue in LT-HSCs and MUTZ-3 cells, and diff_of_classes for TCGA analyses. Genes from CCLE data were pre-ranked by determining mean expression for each gene in AML-high and AML-low cohorts and calculating log_2_ fold change. GSEA was performed using 1000 permutations to determine significance.

### Construction of HSC specific regulatory network

*Cis*-regulatory elements (*cis*REs) govern gene expression via functional interaction with gene promoter directly or indirectly mediated by other *cis*REs^79–81^. To decipher the transcriptional regulation underlying human hematopoiesis, we developed a computational approach called HemeMap, by leveraging a set of multi-omic data in different hematopoietic populations to define *cis*REs, their target genes and their putative regulatory activity throughout hematopoiesis.

### Identification of *cisREs*

To identify the putative *cis*REs in the human hematopoiesis, we used a consensus peak set of ATAC-seq data across 18 cell types across the hematopoiesis, similar to that which we employed in our previous studies^36,43^. The peaks were called using MACS2^82^ for each cell type and uniformly resized to a width of 500 bp centered on the peak summits, then filtered by the ENCODE hg19 blacklist (https://www.encodeproject.org/annotations/ENCSR636HFF/). Peaks uniquely occurring in a particular cell type, i.e. non-overlapping with peaks from other cell types, were retained. For the peaks overlapping in two or more cell types, we compared them iteratively and kept the most significant peak. The remaining peaks were further filtered if they overlapped with gene promoters, which were defined as 500 bp regions around transcription start sites (TSS) of protein coding genes. The *cis*REs from the entire hematopoietic catalog consisted of 432,428 consensus accessible peaks and 18,492 gene promoters.

### Identification of direct interactions

To find the interactions between genes and *cis*REs, we searched for all possible connections between gene and *cis*REs within 500 kb of gene TSS. We used two criteria to define the interactions which the *cis*RE could exert a direct effect on gene regulation: (1) experimental evidence of physical interaction in three-dimensional space or (2) a strong correlation between chromatin accessibility of *cis*RE and target gene expression. To this end, we annotated the nominated links to assess whether *cis*REs and target genes are spatially colocalized (i.e. in a chromatin loop). A published dataset spanning 15 hematopoietic cell types of promoter capture Hi-C (PCHi-C) data was used^42^ and only loops with CHiCAGO score > 5 were considered. Next, we computed ATAC-seq reads falling within *cis*REs across the hematopoietic cell populations and performed normalization using the count per million (CPM) method. We calculated the Pearson correlation coefficient (*r)* between chromatin accessibility of *cis*REs and gene expression across 16 hematopoietic cell types for each possible interaction pair. To determine the significance, we applied Fisher’s *r* to *z* transformation^83^ to correlation coefficients. All the interactions with *r*> 0.345 (equivalent to *P* value < 0.05) were kept. Finally, the nominated links that passed either of these two analyses were retained and a total of 1,218,933 direct interactions were identified.

### Identification of indirect interactions

A gene regulatory network is established by a chain of *cis*REs which connect to the target though direct or indirect manners^79,84^. Previous studies^47,85^ reported that a number of cooperative *cis*REs could associate with the promoter and other *cis*REs related in multi-way contacts in chromatin loops. Co-accessible chromatin has been reported to be highly connected and functionally related^86,87^, which is useful to evaluate the connectivity between *cis*REs. To identify the indirect interactions, we first computed the co-accessibility across 18 cell types between *cis*REs (not including gene promoters) whose genomic distance less than 500 kb. By using the Pearson correlation measurement and Fisher’s *r* to *z* transformation as described above, the co-accessible *cis*RE-*cis*RE links with a correlation coefficient *r* > 0.362 (equivalent to *P* value < 0.05) were selected. Next, to find the shortest path between a *cis*RE and its target promoter, we constructed a regulatory network using the direct gene-*cis*RE interactions and co-accessible *cis*RE-*cis*RE links, and found the shortest paths between *cis*REs and genes in this network. Specifically, the network was built using the igraph R package^88^ with gene-*cis*RE interactions and *cis*RE-*cis*RE links. Dijkstra’s algorithm^89^ is designed for searching for the shortest paths between nodes in a graph. In our network, we used this method to find all the potential indirect interactions mediated by the *cis*REs that have direct gene interactions identified in the first step of our analysis. Given that a smaller weight indicates a greater chance in participating in the shortest path found by the Dijkstra’s method, we added the weight to each edge in the network: weight of a pseudo number of 1e-5 for direct gene-*cis*RE interactions and 1 – *r* for *cis*RE-*cis*RE links, respectively. All of the gene-*cis*RE pairs that did not pass the direct interaction identification were analyzed by Dijkstra’s method. The *cis*REs were filtered out if they were not linked to any gene. In total, 4,315,536 interaction pairs are included in HemeMap.

### HSC specific regulatory network

To define the strengths of *cis*-regulatory interactions in each cell type, we calculated the HemeMap score by using the geometric mean of ATAC-seq signal over all the *cis*REs involved in each interaction to avoid potential bias introduced by the outliers. To get the HSC-specific regulatory network, we used the cumulative Chi-Square distribution to determine an interaction strength threshold of greater than 8.91 which filtered out 95% of the interactions. The remaining interactions were used to build an HSC-specific regulation network containing 12,808 genes and 372,491 *cis*REs.

### De novo motif discovery

To explore the MECOM mediated regulatory network, we retrieved all of the *cis*REs associated with MECOM network genes identified as differentially expressed after *MECOM* editing. We used the 200 bp sequences centered on *cis*REs, i.e. the genomic regions around summits of peaks or TSS, as input for the *de novo* motif discovery analysis. The MEME suite^90^ was used and all the motifs with reported *E* value < 1e-20 were collected from results of DREME^91^ and MEME. Similarity of *de novo* motifs and the putative TF motifs from a comprehensive collection of 401 human TFBS models (HOCOMOCO V11)^92^ was performed using Tomtom^93^. We also correlated the similarity of the ETS family motif identified via *de novo* motif discovery with the EVI1 binding motif from a published dataset^24^ by calculating the Pearson correlation coefficient of the Position Frequency Matrix (PFM) of the two motifs using universal motif R package^94^.

### TF Footprinting analysis

A TF footprint is a particular pattern of Tn5 enzyme cleavage sites generated by ATAC-seq data that enables analysis of chromatin occupancy at the base-pair resolution. There is a depletion of cleavage events at the specific site of TF binding on open chromatin, which allowed for the identification of TF binding events with the consensus motifs of interest from the *de novo* motif discovery analysis^95,96^. For each *de novo* motif, including ETS, RUNX, JUN, KLF, CTCF and GATA, we scanned all of the consensus motif sequences that occur within the *cis*REs in MECOM-mediated regulatory network using the software FIMO^97^ with default parameters, except for a significance threshold of 5E-4. To create a nucleotide resolution cleavage frequency profile for each TF, we used *make_cut_matrix* function (https://github.com/Parkerlab/atactk) to count the Tn5 enzyme cleavage frequency at the recognized motif sites and their flanking +/- 250 bp sequences, using ATAC-seq data from HSCs. Then, we used CENTIPEDE^98^ to build an unsupervised Bayesian mixture model with the cleavage frequency profile to generate a posterior probability value for each motif instance. A motif instance was considered a footprint that is bound by a particular TF when the posterior probability score was greater than 0.95. The plot of cleavage frequency around the footprints was created by aggregating both strands using a custom R script.

### Footprint co-occurrence analysis

To explore how these TFs cooperate with each other via combinatorial binding on the *cis*REs of MECOM network genes, we evaluated the co-occurrence of the TF footprints. Specifically, a hypergeometric test was employed to determine the statistical significance of co-occurrence of two different footprints, as depicted by the following equation:

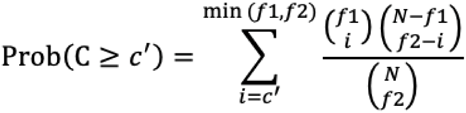

where is the total number of *cis*REs, and are the number of *cis*REs containing footprints of each of the two tested TFs, respectively. *P* value measuring the significance of enrichment is the tail probability of observing or more *cis*REs containing both TF footprints.

### ChIP-seq data analysis

The raw ChIP-seq data^46^ for the binding sites of hematopoietic TFs FLI1, GATA2 and RUNX1 in human CD34+ HSPCs, were downloaded and processed. The paired-end reads were trimmed and aligned to hg19 reference genome using Trimmomatic and Bowtie2, respectively. MACS2^82^ was used for peak calling with the default narrow peak setting. Genomic tracks were generated from BAM files using CPM normalization to facilitate comparison between tracks. The processed CTCF ChIP-seq data from HSPCs and differentiated hematopoietic lineages were obtained from a previous study^52^. To determine the significance of the enrichment of TF occupancy within *cis*REs of MECOM network genes, a permutation test was performed. For each TF, we calculated the number of *cis*REs overlapping with ChIP-seq peaks. The expected distribution of overlapping *cis*REs was generated by 1,000 permutations of an equal number of TF peaks across the genome.. The presence of TF peaks in *cis*REs were counted and the Venn plot was generated by the web app BioVenn^99^. The enrichment of CTCF signal on the footprints was performed using deepTools software^100^. We used Wilcoxon signed-rank test to evaluate the differences of normalized CTCF signals on footprints between HSPCs and other terminal blood cells, namely erythroid cells, T-cells, B-cells, and monocytes.

### CTCF-mediated loop enrichment analysis

A set of 7,358 representative chromatin interactions in hematopoietic cells was identified from a high-resolution Hi-C map of OCI-AML2 cells as previously described^51^. The loops whose anchors overlap with *cis*REs of MECOM down genes were extracted for further analysis. The CTCF-mediated loops (at least one of the anchors containing a CTCF footprint) and non-CTCF-mediated loops (anchors without CTCF footprint) were identified separately. The Low-C data of chromatin looping in LT- and ST-HSC^51^ were normalized by Knight-Ruiz balanced interaction frequencies at a resolution of 25 Kb. We used Juicer to perform aggregate peak analysis (APA)^47^ to test for enrichment of loops within the Low-C data from LT-HSCs and ST-HSCs. Loops containing genes were identified by the genes within the genomic domains between loop anchors. A published RNA-seq data set of CTCF knockdown in LT-HSC was obtained^51^ and we examined the expression of MECOM down gene after CTCF knockdown within CTCF-mediated loops and non-CTCF-mediated loops, respectively. The Wilcoxon signed-rank test was performed to determine the significance.

## ANALYSIS OF PRIMARY AML PATIENT DATA

### Included studies

Three study cohorts were included in the survival analyses. We downloaded RNASeq V2 expression data and corresponding clinical outcomes from the TCGA LAML cohort from cBioPortal (https://www.cbioportal.org/study/summary?id=\laml_tcga_pub)101 for 173 AML patients. The same was done for the BEAT-AML cohort for 430 patients (https://www.cbioportal.org/study/summary?id=\aml_ohsu_2018)54. In addition, the TARGET dataset was downloaded for 440 pediatric AML patients (https://www.cbioportal.org/study/summary?id=\aml_target_2018_pub)55. To gain maximal insight, adult datasets (TCGA and BEAT) were combined, with subsequent adjustments in analyses to account for study specific features. The only pediatric data used was from the TARGET dataset. The results published here are in part based upon data generated by the Therapeutically Applicable Research to Generate Effective Treatments (https://ocg.cancer.gov/programs/target) initiative, phs000218. The data used for this analysis are available at https://portal.gdc.cancer.gov/projects.

### Derivation of variables of interest

We log_2_-transformed the TCGA normalized read counts and stratified the cohort based on MECOM expression (MECOM low, log2(RPKM+1)<4; MECOM high, log2(RPKM+1)≥4). LSC17 score was calculated as follows: (*DNMT3B* × 0.0874) + (*ZBTB46* × −0.0347) + (*NYNRIN* × 0.00865) + (*ARHGAP22* × −0.0138) + (*LAPTM4B* × 0.00582) + (*MMRN1* × 0.0258) + (*DPYSL3* × 0.0284) + (*KIAA0125* × 0.0196) + (*CDK6* × −0.0704) + (*CPXM1* × −0.0258) + (*SOCS2* × 0.0271) + (*SMIM24* × −0.0226) + (*EMP1* × 0.0146) + (*NGFRAP1* × 0.0465) + (*CD34* × 0.0338) + (*AKR1C3* × −0.0402) + (*GPR56* × 0.0501)^58^. For each of the three included studies, the expression of each gene in each individual sample was compared to the mean expression in the pertaining study cohort. GSEA (as described previously) was performed to determine the enrichment or depletion of MECOM down genes in each sample compared to the mean. A sample was determined to have enrichment of MECOM down genes if the Normalized Enrichment Score >0 and p-value <0.05, depletion of MECOM down genes if NES <0 and p-value <0.05, or unchanged MECOM down genes if p-value >0.05. In addition, the normalized enrichment score was studied as a continuous measure of MECOM network status. Clinical risk scoring was provided in tables by each of the studies based on the National Comprehensive Cancer Network criteria, and in this analysis are labelled as Adverse, Intermediate and Favorable for consistency.

### Survival analyses

Kaplan-Meier (KM) curves were constructed demonstrating survival for each cohort (adult and pediatric), and variables (MECOM expression, MECOM network enrichment score, MECOM network enrichment (categorical), LSC17, and clinical risk score). For continuous variables, to appreciate survival differences in the variable in this way, KM curves were stratified by thresholding on the optimum threshold determined by Youden’s J statistic, maximizing both sensitivity and specificity of the metric. Follow-up time was truncated at 2500 days for the pediatric cohort (thereby including n=350, 79.5% of all complete cases), and at 1500 days for the adult cohort (thereby including n=513, 83.8% of all complete cases) for this and subsequent analyses to limit the issue of data sparsity at very late event time points. KM curves were constructed in R using survival and ggsurvplot packages.

Hazard ratios and 95%CI of death were determined from Cox proportional hazards models. These were created for each variable, correcting for contributing study in the adult group. This allowed assessment of continuous variables at their full spectrum. This also allowed for assessment of association of MECOM down network enrichment with mortality, independent of existing clinical approaches such as the clinical risk score and LSC17. Corrected models for age and sex were created and marginal hazard of mortality was derived and displayed graphically by different ages. The R packages’ coxph, survival, rms, ggeffects were used.

For analysis of AML cells from the CCLE database, we downloaded RNASeq and CRISPR dependency data from the Cancer Dependency Map (https://depmap.org)^102–104^. We stratified the cohort based on MECOM expression (MECOM low, log2(RPKM+1)<1; MECOM high, log2(RPKM+1)≥1). Differential essentiality was determined by subtracting the CERES gene effect score of MECOM high-MECOM low AML samples. A negative value indicates stronger essentiality in MECOM-high AML.

## STATISTICAL ANALYSIS

We used unpaired Student’s t-tests for in vitro and in vivo assays of HSC function following MECOM editing (**Fig. 1c,f,h,j, 3b-e,g,I, 5i, 7c,e,h, Extended Data Fig. 1f,g,i,l,p, Extended Data Fig. 3d, Extended Data Fig. 7g**). We used a hypergeometric test to determine the significance of TF footprint co-occurrence (**Figures 4e, 5b, Extended Data Fig. 4f,g**). We used Wilcoxson signed-rank test to determine significance of ETS motifs in cisREs of MECOM network genes (**Extended Data Fig. 4e**), differential CTCF binding during hematopoiesis (**Extended Data Fig. 5a**), differential rescue of MECOM down genes with CTCF footprints after concurrent *CTCF* perturbation (**Extended Data Fig. 5j,k**). We used the Mantel-Cox log-rank test for analyzing survival (**Figures 6c-g, and Extended Data Fig. 6f-j**). We used the Chi-squared test to determine the significance of HemeMap scores (**Extended Data Fig. 4d**). The significance of motif discovery was calculated using Fisher’s exact test (**Fig. 4b**). We used Pearson correlation and Fisher’s to transformation to determine significant interactions in the establishment of HemeMap (**Fig. 4a**). Pearson correlation is also used to compare the ETS motif in our analysis to the EVI1 binding motif (**Fig. 4c**). The Kolmogorov Smirnov (K-S) test was used to determine the significance of GSEA (**Fig. 3h,j, 5j,k, 7f,g, Extended Data Fig. 3f,g, Extended Data Fig. 5h,i).** Mann-Whitney U test was used in the 10x scRNA-seq analysis performed with Pegasus 1.0. Statistical analyses were performed using Prism v8.4, the R (version 3.6.3) language for Statistical Computing, and Python (version 3.7.7). Parameters such as sample size, number of replicates, measures of center, dispersion, precision (mean ± s.e.m) and statistical significance are reported in the Figures and Figure Legends. All measurements were taken from distinct samples unless otherwise noted.

## DATA AND CODE AVAILABILITY

Code and source data for reproducing results of this study are available on GitHub (https://github.com/sankaranlab/mecom_var).

