## Supplementary figures and images for "A genetic disorder reveals a hematopoietic stem cell regulatory network co-opted in leukemia"

### Extended Data Fig. 1

#### Extended Data Figure 1

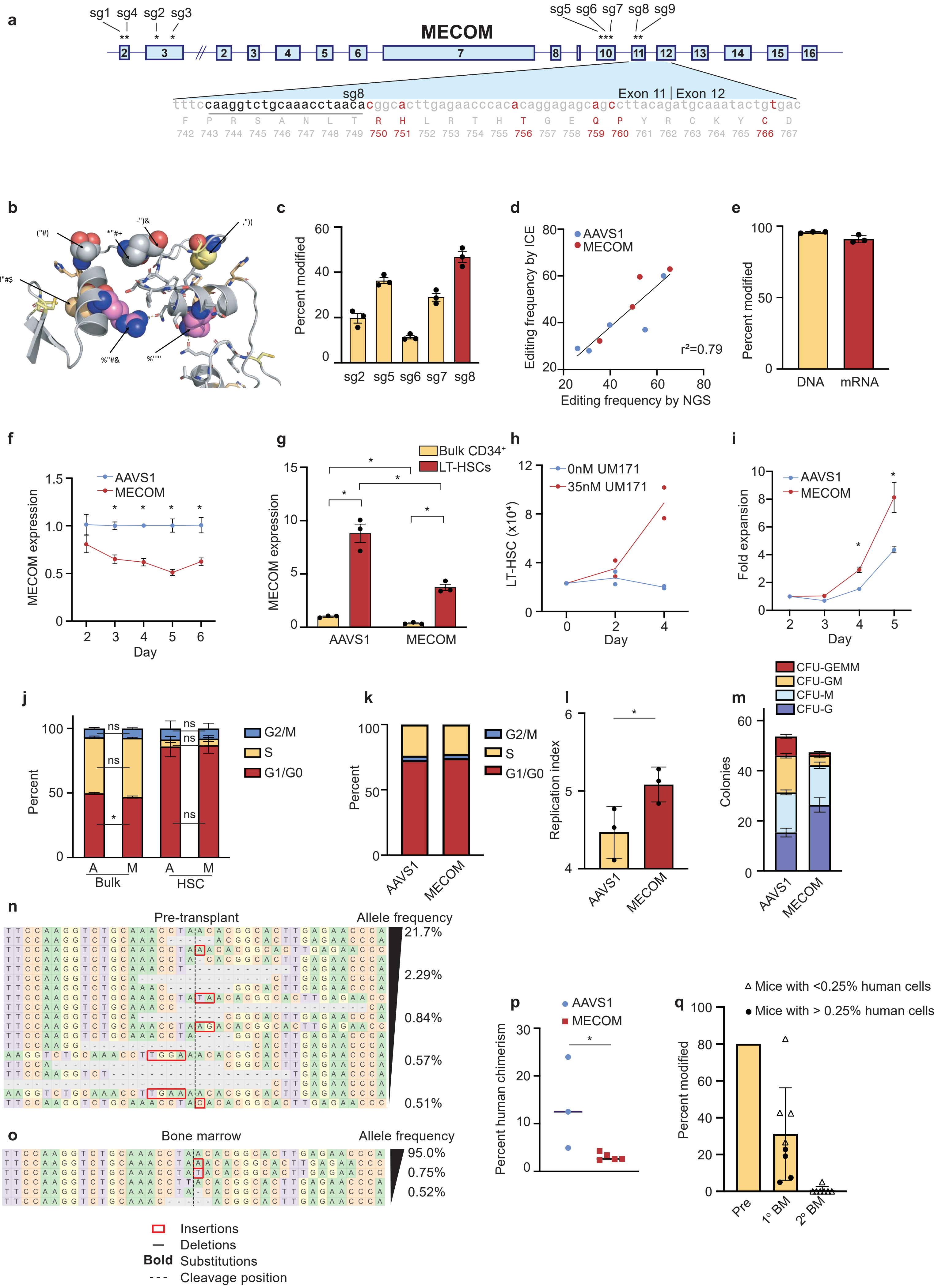

### Extended Data Fig. 2

**Extended Data Figure 2**

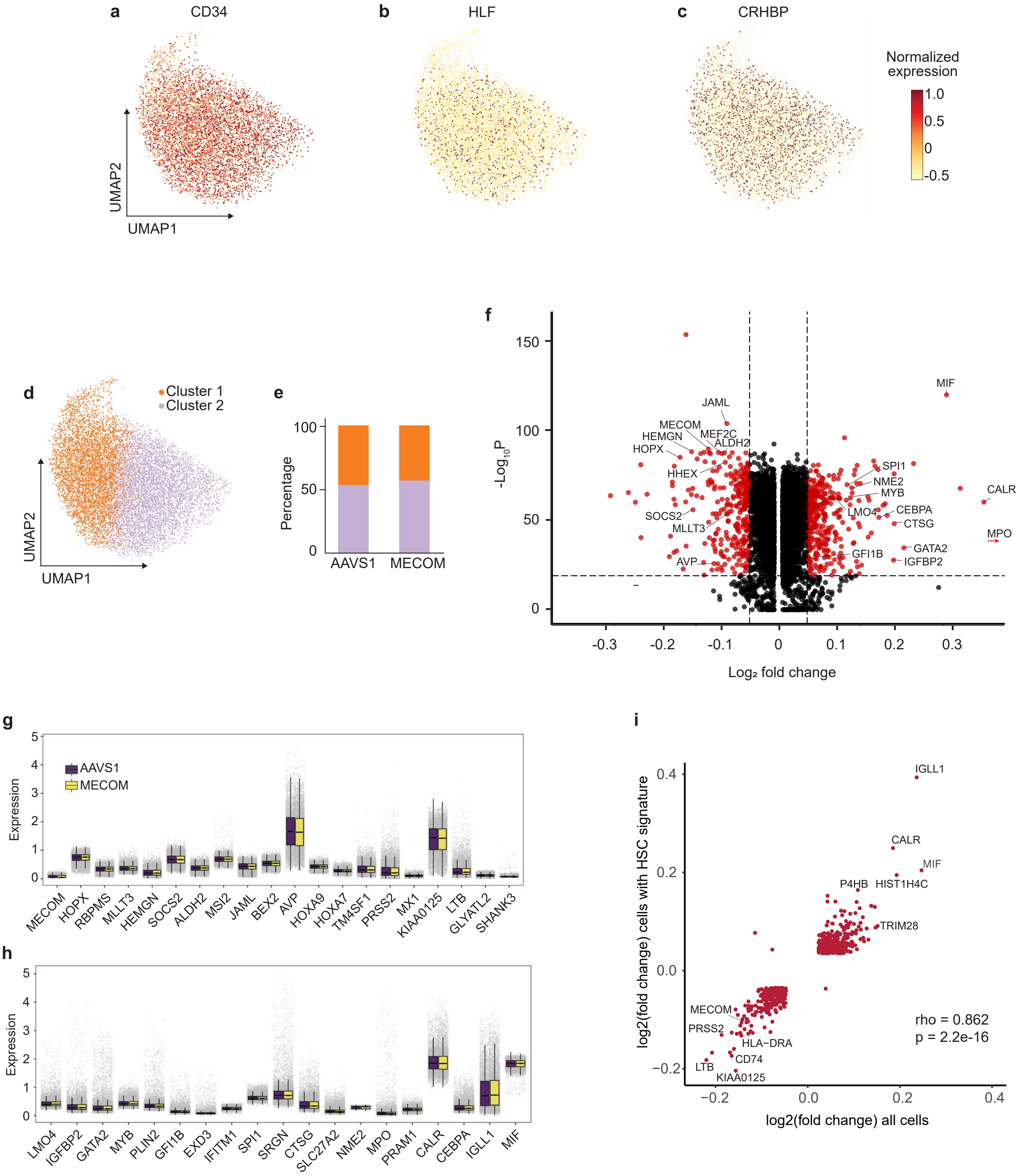

### Extended Data Fig. 3

#### Extended Data Figure 3

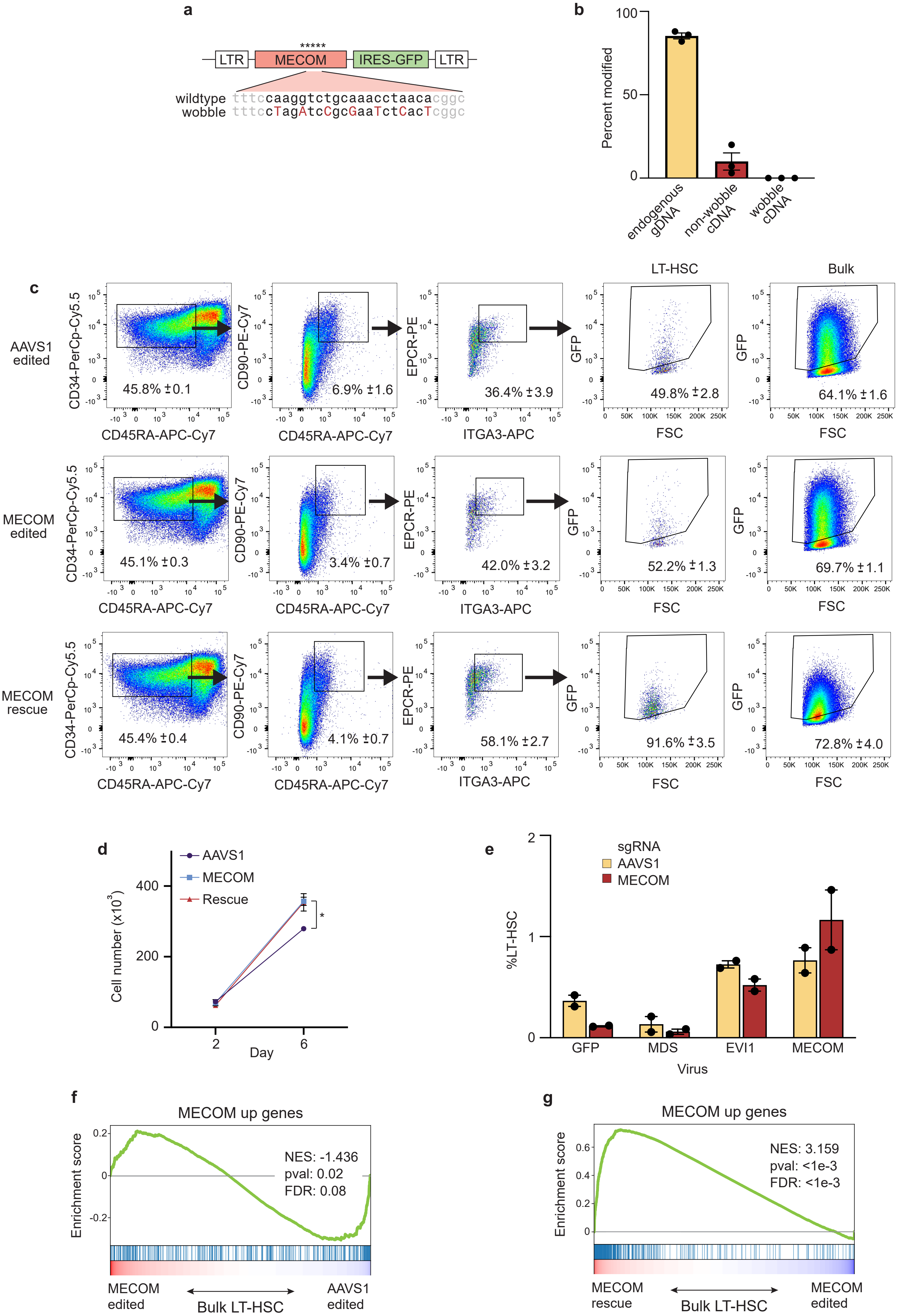

### Extended Data Fig. 4

**Extended Data Figure 4**

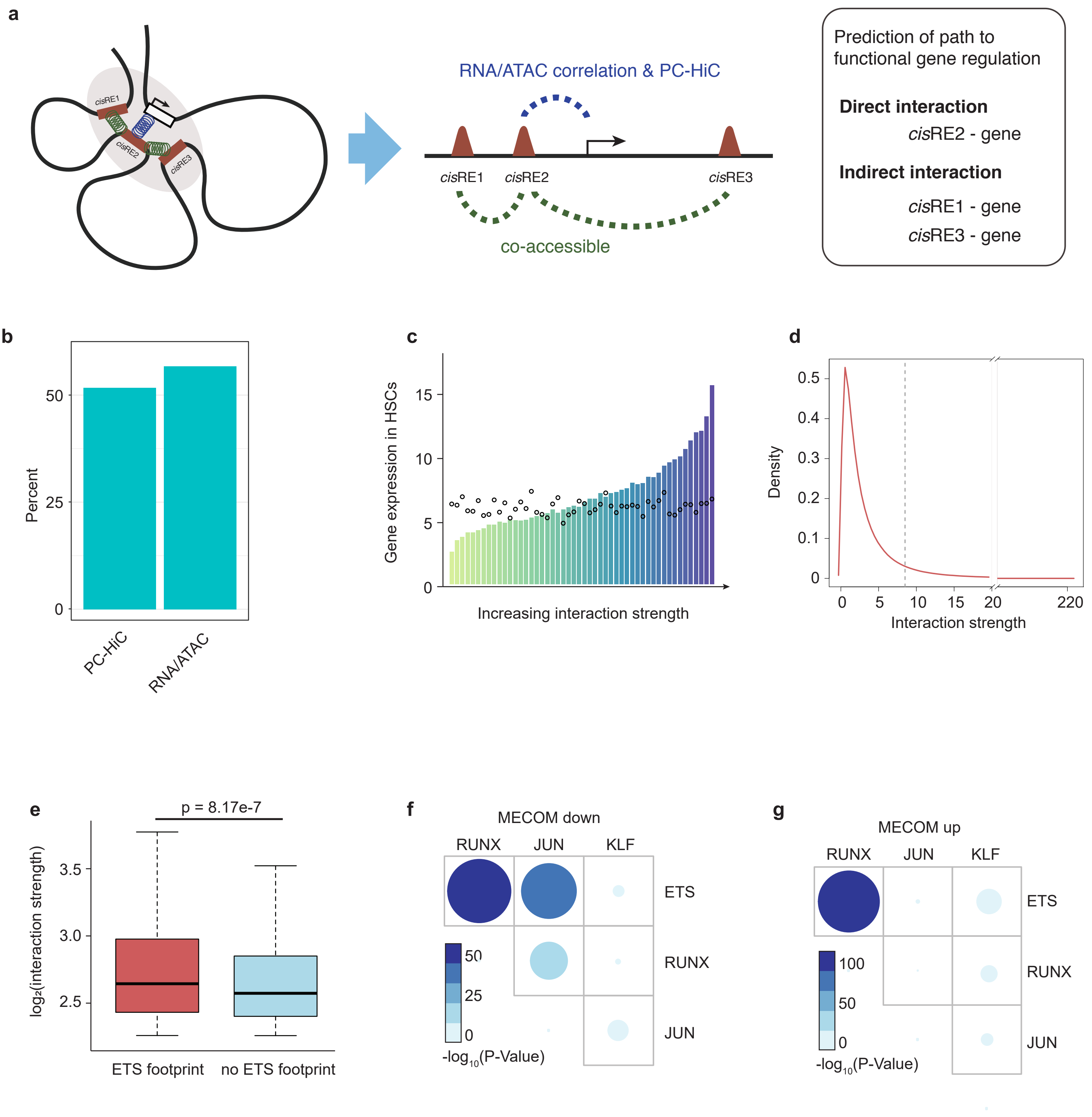

### Extended Data Fig. 5

**Extended Data Figure 5**

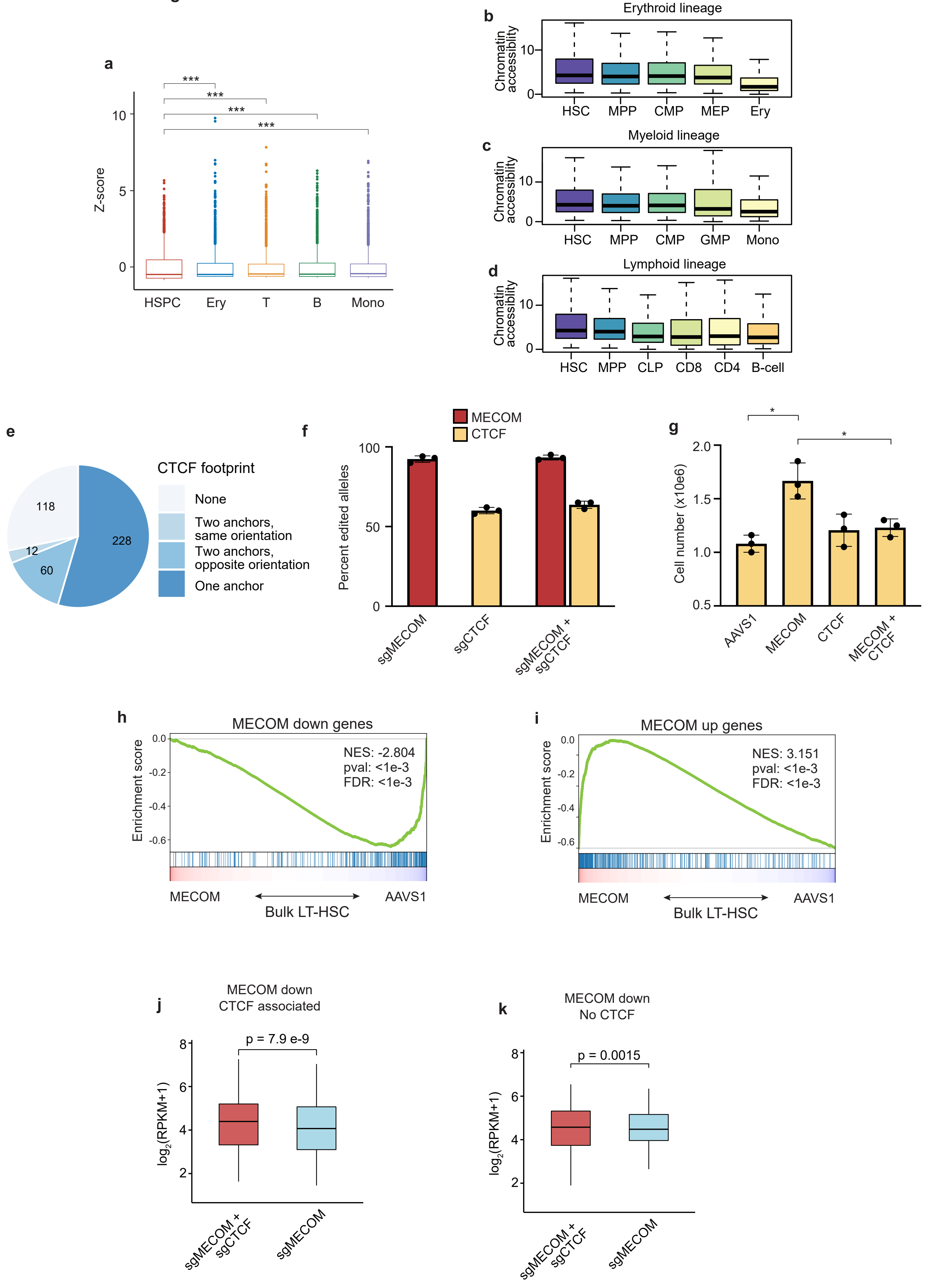

### Extended Data Fig. 6

**Extended Data Figure 6**

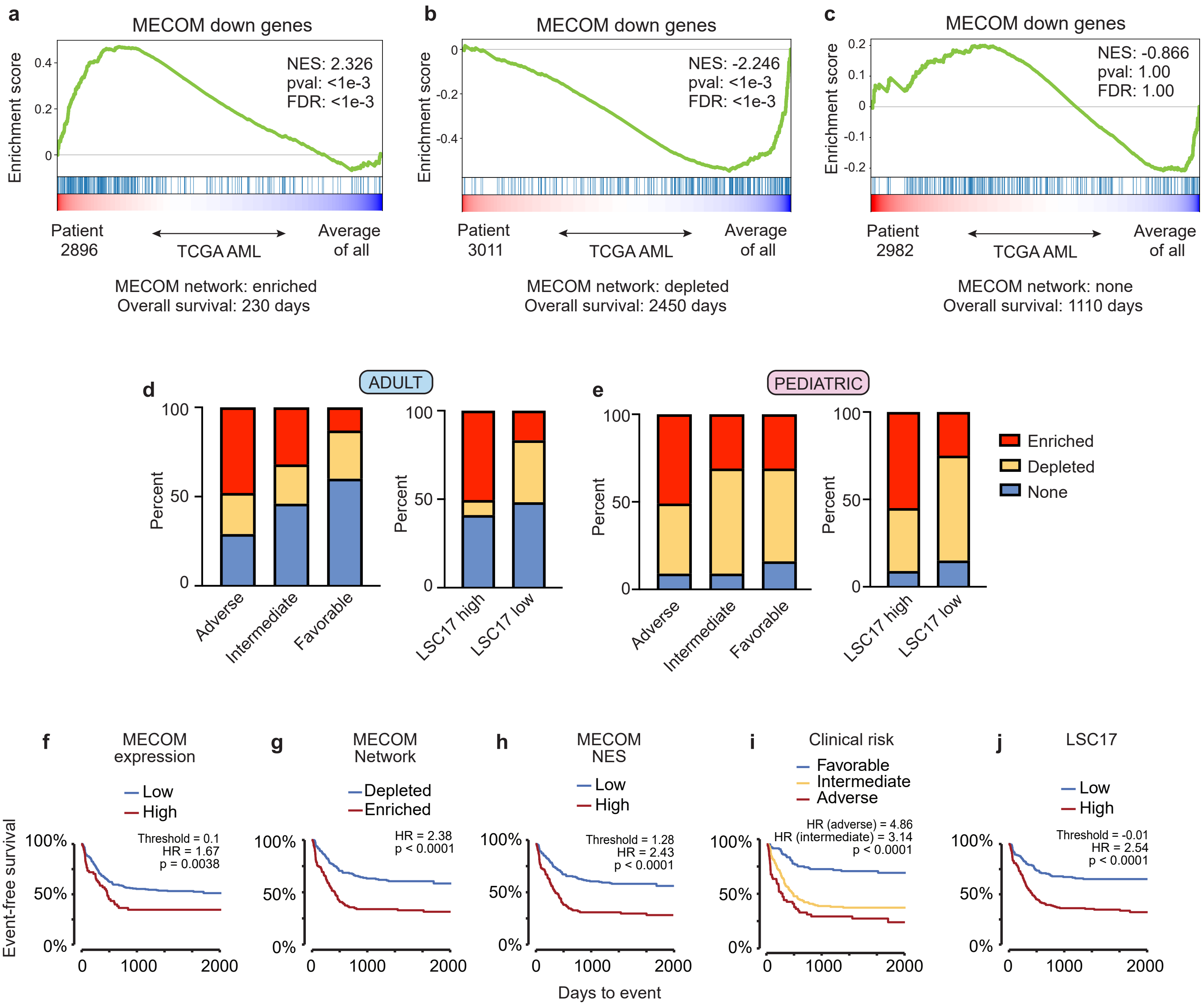

### Extended Data Fig. 7

**Extended Data Figure 7**

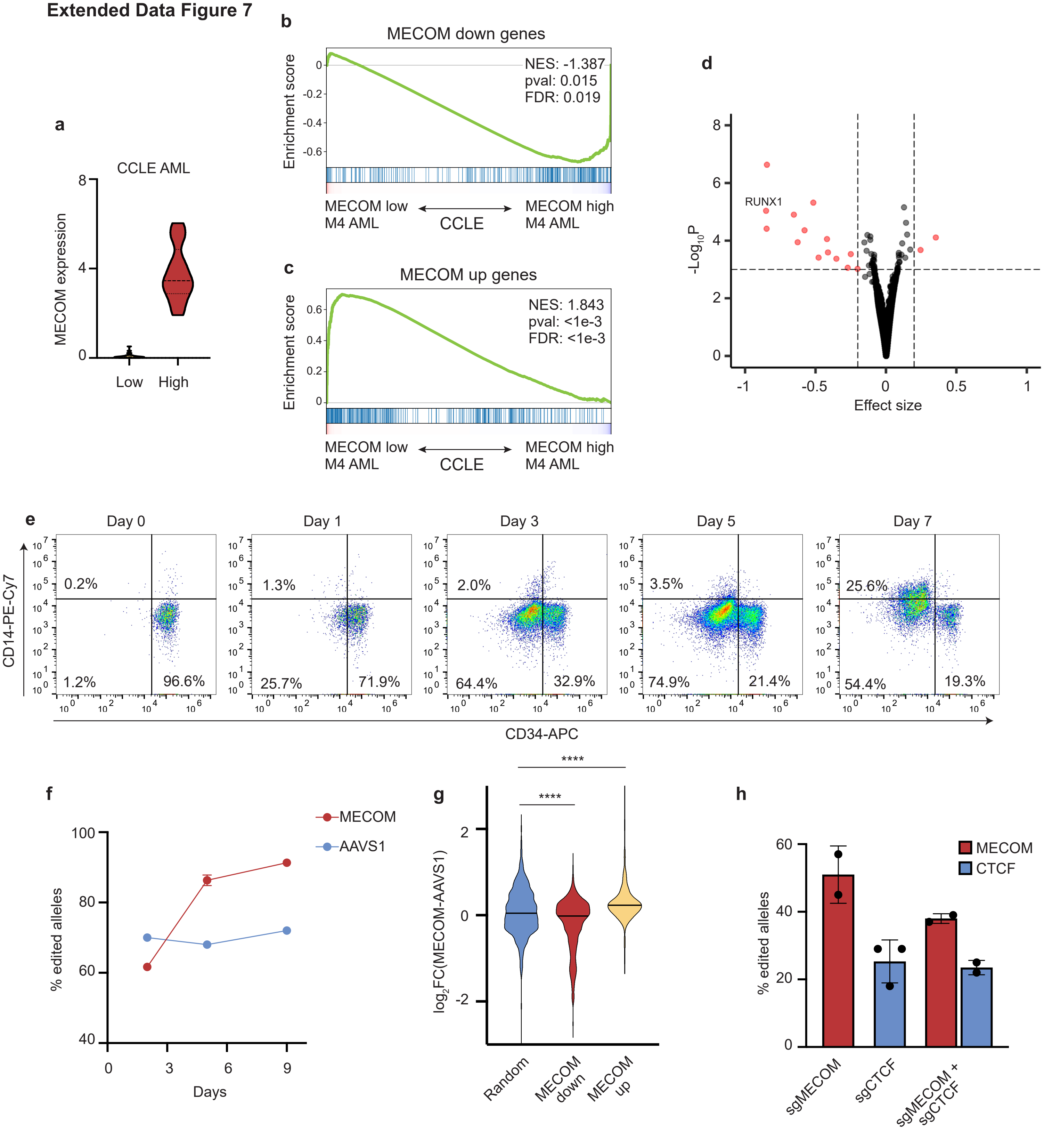
